# Collateral lethality between *HDAC1* and *HDAC2* exploits cancer-specific NuRD complex vulnerabilities

**DOI:** 10.1101/2022.05.30.493851

**Authors:** Yuxiang Zhang, David Remillard, Ugoma Onubogu, Barbara Karakyriakou, Joshua N. Asiaban, Anissa R. Ramos, Kirsten Bowland, Timothy R. Bishop, Christopher J. Ott, Michalina Janiszewska, Benjamin F. Cravatt, Michael A. Erb

**Affiliations:** Department of Chemistry, The Scripps Research Institute, La Jolla, California, USA; Department of Molecular Medicine, UF Scripps Biomedical Research, Jupiter, Florida, USA; Massachusetts General Hospital Cancer Center, Charlestown, Massachusetts, USA; Department of Medicine, Harvard Medical School, Boston, Massachusetts, USA; Broad Institute of MIT & Harvard, Cambridge, Massachusetts, USA

## Abstract

Histone deacetylases (HDACs) have been widely pursued as targets for anti-cancer therapeutics. However, many of these targets are universally essential for cell survival, which may limit the therapeutic window that can be achieved by drug candidates. By examining large collections of CRISPR/Cas9-based essentiality screens, we discovered a genetic interaction between *HDAC1* and *HDAC2* wherein each paralog is synthetically lethal with hemizygous deletion of the other. This collateral synthetic lethality is caused by recurrent chromosomal translocations that occur in diverse solid and hematological malignancies, including neuroblastoma and multiple myeloma. Using genetic deletion or dTAG-mediated degradation, we show that HDAC2 disruption suppresses the growth of *HDAC1*-deficient neuroblastoma *in vitro* and *in vivo.* Mechanistically, we find that targeted degradation of HDAC2 in these cells prompts the degradation of several members of the nucleosome remodeling and deacetylase (NuRD) complex, leading to diminished chromatin accessibility at HDAC2/NuRD-bound sites of the genome and impaired control of enhancer-associated transcription. Furthermore, we reveal that several of the degraded NuRD complex subunits are dependencies in neuroblastoma and multiple myeloma, providing motivation to develop paralog-selective HDAC1 or HDAC2 degraders. Altogether, we identify *HDAC1/2* collateral synthetic lethality as a new therapeutic target and reveal a novel mechanism for exploiting NuRD-associated cancer dependencies.

## Introduction

Currently, 4 HDAC inhibitors are approved for use as anti-cancer agents in humans^1^, but the vast majority of clinical development efforts have failed due to low response rates and poor tolerability^2^. Notably, all but one of the currently approved HDAC inhibitors are broadly active against most or all HDAC enzymes, which likely contributes to the toxicities that have been observed in clinical trials^2^. This has prompted broad efforts to develop class-specific HDAC therapeutics, like romidepsin (FK228), which targets class I HDACs and is approved for the treatment of cutaneous T cell lymphoma and peripheral T cell lymphoma. Class I HDACs include HDAC1, HDAC2, HDAC3 and HDAC8. These enzymes are predominantly localized to the nucleus and catalyze zinc-dependent removal of acyl groups from the ε-amino lysine of various histone and non-histone substrates^3^. HDAC1 and HDAC2, which were originally discovered through target identification studies for the HDAC inhibitor, trapoxin^4^, are present in multiple protein complexes, including NuRD, Sin3A, CoREST, MiDAC, and SMRT/NCoR^3,5^. These paralogs regulate diverse cellular processes through their transcriptional co-regulatory function and play key roles in normal development as well as tumorigenesis^3,6^. Unfortunately, multiple recent studies have revealed that simultaneous genetic disruption of *HDAC1* and *HDAC2* is pan-lethal^7,8^, explaining, at least in part, the difficulty of achieving a therapeutic window in patients, even with Class-I-selective HDAC inhibitors.

In cancer biology, synthetic lethality is used as a framework to identify genes that are only essential for cell survival in the presence of particular tumor-associated alterations^9^. As drug targets, these gene products promise a favorable therapeutic window since many or most normal cell types do not require them for survival. The study of synthetic lethality has historically depended on hypothesis-driven research that is well demonstrated by the classic example of PARP inhibition in *BRCA1/2*-mutated cancers^10^. However, the increasingly sophisticated development of genetic loss-of-function screening tools for mammalian cells has enabled the unbiased discovery of many additional examples, supporting new therapeutic hypotheses for drug discovery. Pooled genetic screens have been particularly informative, revealing, for example, synthetic lethality between *SMARCA2* and *SMARCA4^11–13^, CBP* and *p300^14^,* and *STAG1* and *STAG2*^15^. To that end, the pre-publication release of large-scale CRISPR screening datasets, well-exemplified by the Cancer Dependency Map from the Broad Institute^16,17^, has already proven exceptionally useful for new target discovery.

It is worth noting that paralogous proteins with redundant or semi-redundant function represent a major subset of the synthetic lethal pairs identified to date^9^. Paralogs commonly share structural similarity and functional redundancy, which can make the inactivation of one gene inconsequential for cell survival. However, one paralog can become indispensable when loss-of-function alterations compromise the other. Therefore, surveying paralogous proteins by combinatorial genetic screens, or examining existing dependency maps for genetic interactions between paralogs, could expedite the identification of actionable synthetic lethalities in cancer^7,18,19^.

In this study, we scrutinized the Cancer Dependency Map to examine the requirement for individual Class I HDACs across dozens of human cancers. Through this effort, we identified a previously unrecognized collateral synthetic lethality between *HDAC1* and *HDAC2* caused by recurrent cancer-associated chromosomal deletions. *HDAC1* is located in proximity to a region of chromosome 1p that is deleted in approximately 20% of neuroblastoma^20–23^. Likewise, *HDAC2,* is located within chromosome 6q, which experiences deletions in nearly 40% of lymphoid malignancies^24–27^. We demonstrate that hemizygous deletion of either gene sensitizes cancer cells to loss of the other paralog, owing to the fact that combined HDAC1/2 function is pan-essential. Using complementary genetic and pharmacological approaches, we investigated the effects of targeting this synthetic lethality and identified an unexpected link between HDAC1/2 abundance and destabilization of the NuRD chromatin remodeler complex. Altogether, these data motivate the discovery of paralog selective HDAC1 or HDAC2 degraders for the development of new anti-cancer therapeutics.

## Results

To systematically evaluate the requirement for individual class I HDACs in cancer, we extracted scaled gene-level dependency scores (Chronos) from the 21Q4 release of the Cancer Dependency Map (DepMap) and calculated the average effect of disrupting each HDAC across 25 cancer lineages **(Fig. 1a)**. In contrast to *HDAC3,*which is indiscriminately required for cell survival, most cancer lineages are not affected by individual deletions of *HDAC1* or *HDAC2* **(Fig. 1a)**. However, we noticed that multiple myeloma cell lines categorized under the plasma cell lineage are highly dependent on *HDAC1,* whereas neuroblastoma cell lines categorized under the peripheral nervous system are highly dependent on *HDAC2* **(Fig. 1a)**. To contextualize these effects, we compared dependency scores between neuroblastoma and all other lineages for 18,119 genes included in the cancer DepMap **(Fig. 1b)**. *HDAC2* registers as one of the most selective dependencies in neuroblastoma and is comparable to many of the core regulatory circuitry (CRC) transcription factors (TF) that are known to be selectively required by neuroblastoma cells^28,29^ **(Fig. 1b and Extended Data Fig. 1a)**. These data are consistent with the nomination of *HDAC2* as a neuroblastoma dependency by the Pediatric Cancer DepMap^30,31^. Likewise, a significantly higher dependency on *HDAC1* is observed in multiple myeloma compared to other cancer types, although the effect was not as selective as *HDAC2* in neuroblastoma **(Fig. 1c)**. These data demonstrate that *HDAC1* and *HDAC2* are not ordinarily essential genes but can be required under certain genetic contexts.

**Figure 1:**
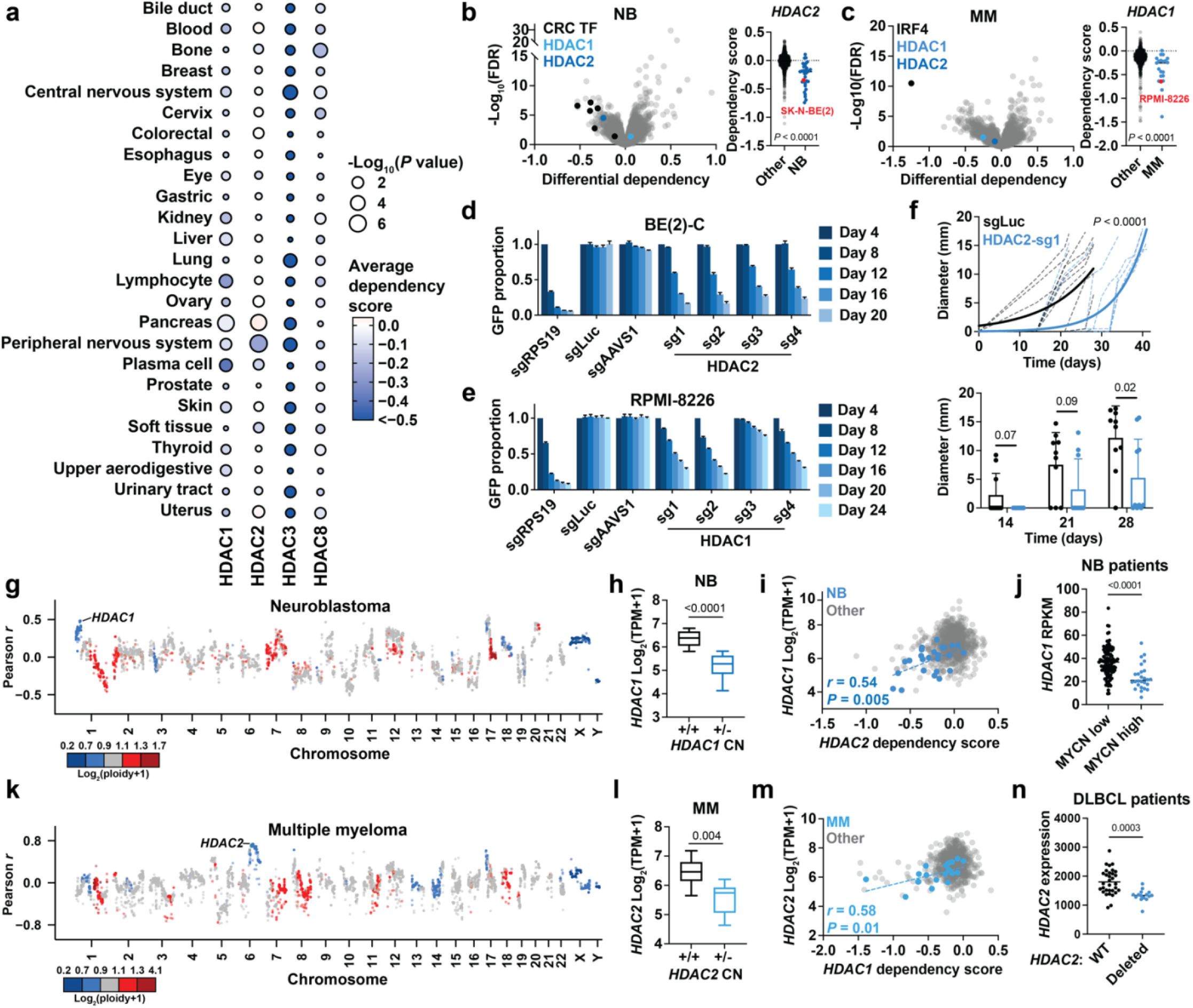
Collateral synthetic lethality between *HDAC1* and *HDAC2.* **a**, Average Chronos gene effect scores (dependency scores) shown for each cancer lineage (circle color). *P* value (circle size) given by two-tailed Student’s *t*-test comparing cell lines within each lineage to all other cell lines. **b**, Left, volcano plot of differential dependencies (average dependency score in neuroblastoma compared to average of all other cell lines). False discovery rate (FDR) determined by *P* values given by two-tailed Student’s *t*-test. Right, dependency scores of *HDAC2* in neuroblastoma lines (blue, *n* = 34) and all other lines (black, *n* = 1,020). **c**, Left, volcano plot of differential dependencies in multiple myeloma. Right, dependency scores of *HDAC1* in multiple myeloma cell lines (blue, *n* = 21) and all other lines (black, *n* = 1,033). **d,** CRISPR-Cas9-based competitive growth assays in BE(2)-C cells. Negative controls include nontargeting (sgLuc) and cutting (sgAAVS1). All measurements of GFP-positive proportion are normalized to the day 4 measurements in the same group. Mean ± s.e.m., *n* = 3. **e,** As in (**d**) for RPMI-8226 cells. **f**, Top, growth of individual subcutaneous BE(2)-C-Cas9 xenografts expressing sgLuc or HDAC2-sg1 (*n* = 10 per group). Non-linear fit with exponential (Malthusian) model shown for each group (solid lines). *P*value determined by the extra-sum-of-squares F test. Bottom, tumor diameters over time. Mean ± s.d., *n* = 10. **g**, Correlations between gene copy number and *HDAC2* dependency in neuroblastoma cell lines on DepMap (Pearson, *n* = 34 neuroblastoma cell lines). Points colored by average copy number in neuroblastoma cell lines. **h**, Boxplot of *HDAC1* expression in neuroblastoma cell lines with (*n* = 12) and without (*n* = 19) *HDAC1* deletion (see **Methods** for cutoff). **i**, Correlation of *HDAC2* dependency versus *HDAC1* expression in neuroblastoma cell lines and other lineages (*n* = 973). *P*value was determined by Pearson correlation coefficient (*r*). **j**, *HDAC1* expression in neuroblastoma patient sample with low *MYCN* expression (black, *n* = 115) and high *MYCN* expression (black, *n* = 26) (cutoff: RPKM < 90) (see **Methods** for data accessibility). **k**, Correlation between gene copy number and *HDAC1* dependency in multiple myeloma (*n* = 21). **l**, Boxplot of *HDAC2* expression in multiple myeloma lines with (*n* = 10) and without (*n* = 20) *HDAC2* deletion (see **Methods** for cutoff). **m**, *HDAC1* dependency versus *HDAC2* expression in multiple myeloma and other lineages (*n* = 973). *P* value was determined by Pearson correlation coefficient (*r*). **n**, *HDAC2* expression in DLBCL patients with wild-type *HDAC2* (black, *n* = 32) and *HDAC2* loss (blue, *n* = 15) (see **Methods** for data accessibility). Unless specified, *P* values were determined by two-tailed Student’s *t*-test. Boxplots represent 25-75 percentiles with whiskers extending to 10-90 percentiles.

To validate *HDAC1* and *HDAC2* as dependencies in these cancer subtypes, we performed CRISPR-Cas9-based competitive growth assays using four independent sgRNAs targeting *HDAC1* or *HDAC2* **(Extended Data Fig. 1b)**. Each was confirmed to mediate on-target effects by immunoblot analysis and a negative control sgRNA targeting the *AAVS1* locus was validated by TIDE (tracking of indels by decomposition)^32^ **(Extended Data Fig. 1c,d)**. We found that *HDAC2* disruption impairs the competitive proliferation of BE(2)-C neuroblastoma cells, whose parental line, SK-N-BE(2), shows a requirement for *HDAC2* on DepMap **(Fig. 1b,d)**. Meanwhile, *HDAC1* disruption impairs the competitive proliferation of RPMI-8226 cells, which are among the more sensitive multiple myeloma cell lines on DepMap **(Fig. 1c,e)**. We extended these findings *in vivo* using a subcutaneous xenograft model of neuroblastoma wherein we observed delayed tumor growth as a result of *HDAC2* disruption by CRISPR/Cas9 **(Fig. 1f)**.

By integrating copy number information from the Cancer Cell Line Encyclopedia (CCLE)^33^ with *HDAC2* dependency scores in neuroblastoma, we discovered that sensitivity to *HDAC2* disruption correlates with chromosome 1p deletions that encompass the *HDAC1* locus **(Fig. 1g)**. This is consistent with our demonstration that BE(2)-C neuroblastoma cells, which harbor a monoallelic loss of *HDAC1***(Extended Data Fig. 2a)**, are sensitive to loss of *HDAC2* **(Fig. 1d)**. We therefore hypothesized that deletion of *HDAC1* creates a synthetic vulnerability to HDAC2 disruption by reducing *HDAC1* expression. In support of this hypothesis, we find that *HDAC1* expression is both significantly lower in neuroblastoma cells with an *HDAC1* deletion and also correlated with *HDAC2* dependency **(Fig. 1h,i)**. Importantly, we note that the *HDAC1* locus (1p35) is proximal to 1p36, which is deleted in nearly a quarter of neuroblastoma cases, particularly those with *MYCN* amplifications, a key feature of aggressive disease^20,23^. In CCLE, we observe *HDAC1* deletions in 12 of 31 neuroblastoma cell lines (39%, **Extended Data Fig. 2a)**, suggesting *HDAC1* deletions are a frequent consequence of 1p36 deletions. Furthermore, we analyzed neuroblastoma patient data available on cBioPortal^34^ and found that *MYCN*-amplified tumors feature significantly reduced *HDAC1* expression **(Fig. 1j)**. This confirms that collateral deletions of *HDAC1,* which sensitize cells to *HDAC2* disruption, are a common feature of high-risk, *MYCN*-amplified neuroblastoma.

**Figure 2:**
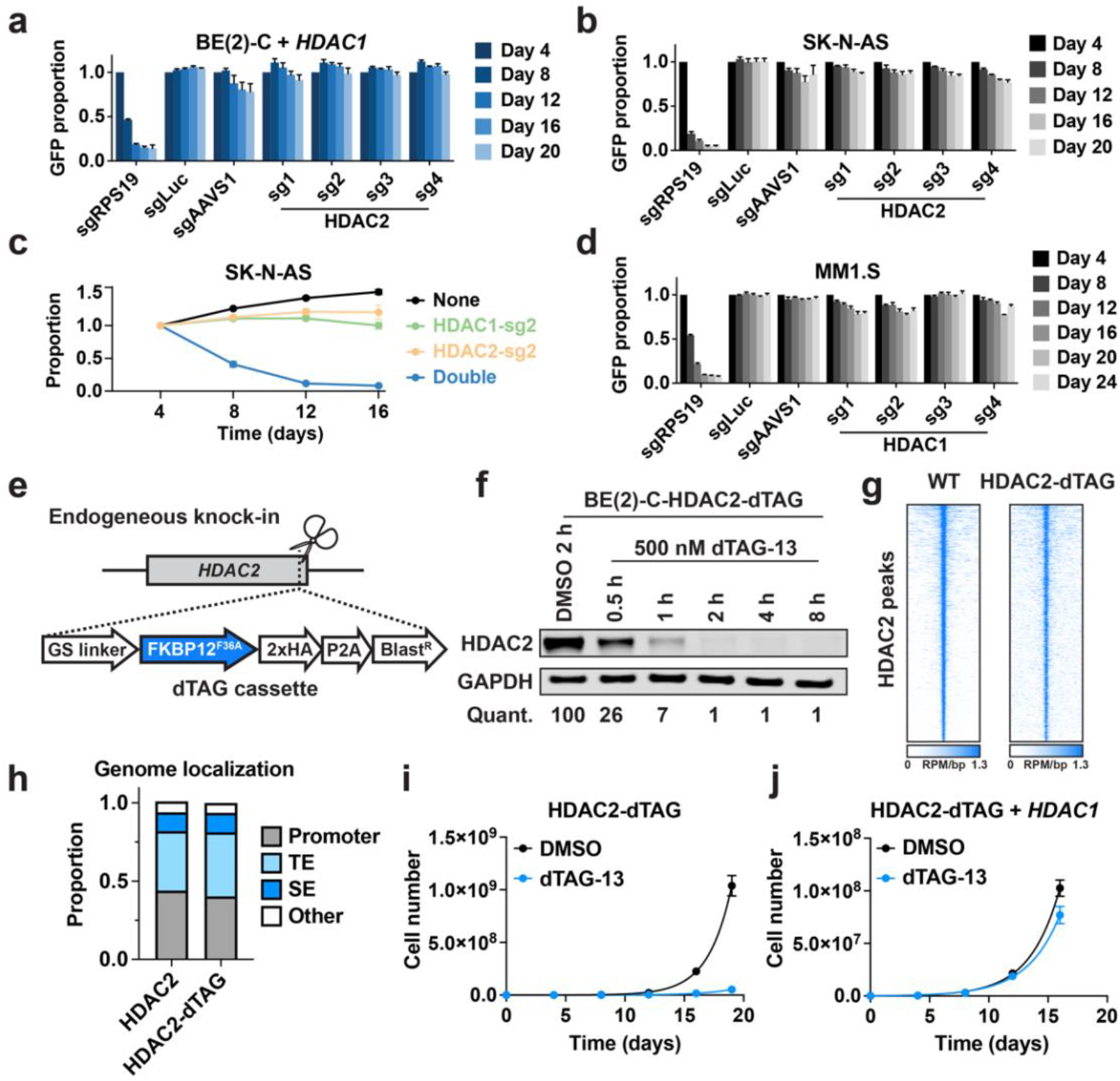
Hemizygous *HDAC1* deletion is necessary and sufficient to sensitize neuroblastoma to loss of HDAC2. **a**, Competitive growth assay in BE(2)-C cells overexpressing *HDAC1.* Mean ± s.e.m., *n* = 3. **b,**Competitive growth assay in SK-N-AS cells. Mean ± s.e.m., *n* = 3. **c**, Two-color competitive growth assay in SK-N-AS cells. Mean ± s.d., *n* = 3. Proportion of each sub-population normalized to day 4. **d**, Competitive growth assay in MM1 .S cells. Mean ± s.e.m., *n* = 3. **e**, Schematic depiction of dTAG knock-in. **f**, Kinetics of HDAC2-dTAG degradation in BE(2)-C-HDAC2-dTAG cells following dTAG-13 treatment (500 nM). DMSO-normalized quantification shown below. **g**, Rank-ordered heatmaps of CUT&RUN signal for HDAC2 in wild-type BE(2)-C cells and HA in BE(2)-C-HDAC2-dTAG cells (ranked based on HDAC2 signal at HDAC2 binding sites in wild-type BE(2)-C cells). **h**, Genomic feature distribution of HDAC2- and HDAC2-dTAG-bound sites. Promoters are defined as transcription start site (TSS) ± 1kb. Enhancers are defined by TSS-distal H3K27ac signal by ROSE2. TE, typical enhancer. SE, super enhancer. **i**,**j** Proliferation of BE(2)-C-HDAC2-dTAG cells (**i**) and BE(2)-C-HDAC2-dTAG with *HDAC1* overexpression (**j**) in response to dTAG-13 treatment (500 nM). Mean ± s.e.m., *n* = 3.

Repeating these analyses in multiple myeloma, we discovered that 6q deletions, which cause hemizygous *HDAC2* deletions, correlate with sensitivity to loss of *HDAC1* **(Fig. 1k)**. Moreover, *HDAC2* deletion is associated with reduced *HDAC2* expression **(Fig. 1l)**, which is in turn correlated with sensitivity to *HDAC1* disruption by CRISPR/Cas9 **(Fig. 1m)**. This is consistent with our demonstration that RPMI-8226 multiple myeloma cells, which harbor an *HDAC2* deletion **(Extended Data Fig. 2b)**, are sensitive to loss of *HDAC1* in competitive proliferation assays **(Fig. 1e)**. Since 6q deletions are known to be common in multiple myeloma as well as several other lymphoid malignancies^24–26^, *HDAC1* is likely to represent a broad dependency in multiple cancer subtypes. Indeed, we find that diffuse large B-cell lymphoma (DLBCL) are significantly more dependent on *HDAC1* than other lineages **(Extended Data Fig. 1e)** and have significantly reduced HDAC2 expression in both cell lines and human patient samples with *HDAC2* deletions **(Fig. 1n and Extended Data Fig. 1f)**. Based on these data, we hypothesized that *HDAC1* and *HDAC2* form a reciprocal synthetic lethality caused by hemizygous chromosomal deletions that impact a broad range of solid and hematological malignancies.

To functionally validate that *HDAC1* deletion sensitizes cells to loss of *HDAC2,* we exogenously expressed *HDAC1* in BE(2)-C neuroblastoma cells by lentiviral transduction and found that this fully rescued the anti-proliferative effects of *HDAC2* disruption by CRISPR/Cas9 **(Fig. 2a)**. As further validation, we tested *HDAC2* disruption in SK-N-AS neuroblastoma cells, which do not harbor an *HDAC1* deletion and express higher levels of HDAC1 protein than BE(2)-C cells **(Extended Data Fig. 2a,c)**. Unlike BE(2)-C cells, SK-N-AS cells are insensitive to loss of *HDAC2* **(Fig. 2b)**. We noted that *HDAC1* and *HDAC2* knockdown by RNA interference has previously been performed in KELLY and BE(2)-C cells to understand the pro-differentiation effects of HDAC1/2 inhibition in neuroblastoma^35^. In this study, knockdown of *HDAC2* elicited anti-proliferative effects in BE(2)-C cells but not in KELLY cells, which do not possess an *HDAC1* deletion. However, combined knockdown of both *HDAC1* and *HDAC2* inhibited the growth of both cell lines. Likewise, we demonstrated that combined deletion of *HDAC1* and *HDAC2* dramatically impairs the competitive fitness of SK-N-AS cells, confirming that loss of *HDAC1* can sensitize cells to loss of *HDAC2* **(Fig. 2c and Extended Data Fig. 2d)**. Likewise, in multiple myeloma, we find that *HDAC1* is not required for the growth of MM1.S cells, which do not harbor an *HDAC2* deletion **(Fig. 2d and Extended Data Fig. 2b,c)**. Altogether, these data demonstrate that *HDAC1* or *HDAC2* deletions are necessary and sufficient to sensitize cancer cells to loss of the remaining paralog. Since combined deletion of *HDAC1* and *HDAC2* is toxic to most cell types^7^,^8^ - including, as we reproduce here, to the AML cell line, OCI-AML2 **(Extended Data Fig. 2e)** – we propose that paralog-selective therapeutics would support an improved therapeutic window for drug development than HDAC1/2 inhibitors.

To enable an orthogonal validation of these data, we employed the dTAG system for chemically induced protein degradation. We reasoned that the rapid kinetics of this approach would support subsequent mechanistic investigations while also modeling the effects of therapeutic disruption in a manner that could inform future undertakings in drug discovery^36,37^. The dTAG system relies on proteolysis targeting chimeras (PROTACs) to elicit rapid and selective degradation of proteins that have been tagged with an engineered “bump-and-hole” variant of FKBP12^38,39^ **(Extended Data Fig. 2f)**. Using CRISPR/Cas9, we engineered BE(2)-C cells to express HDAC2 with a carboxy-terminal fusion of FKBP12^F36V^ **(Fig. 2e and Extended Data Fig. 2g,h)**. This “HDAC2-dTAG” chimera can be degraded by the PROTAC, dTAG-13, in a dose-responsive and time-dependent fashion, with virtually complete degradation achieved within 2 hours **(Fig. 2f and Extended Data Fig. 2i)**. To confirm that the tag does not interfere with the normal genomic distribution of HDAC2, we performed Cleavage Under Targets and Release Using Nuclease (CUT&RUN)^40^. These studies revealed a strong correlation between HDAC2-dTAG localization in BE(2)-C-HDAC2-dTAG cells and HDAC2 localization in wild-type BE(2)-C cells **(Fig. 2g and Extended Data Fig. 2j),** with both proteins occupying promoters and H3K27ac-defined typical enhancers (TE) and super enhancers (SE) at similar frequencies **(Fig. 2h)**. HDAC2-dTAG also retains protein-protein interactions with members of the NuRD complex **(Extended Data Fig. 2k)**, altogether validating the dTAG system as a complementary method to study HDAC2 function in BE(2)-C cells. To test whether HDAC2 degradation recapitulates the anti-proliferative effect of *HDAC2* disruption by CRISPR/Cas9, we measured the growth of BE(2)-C and BE(2)-C-HDAC2-dTAG cells in response to dTAG-13. As expected, the growth of wild-type cells was not affected by dTAG-13 **(Extended Data Fig. 2l)**, but it potently inhibited the growth of BE(2)-C-HDAC2-dTAG cells **(Fig. 2i)**. Importantly, overexpression of *HDAC1* was able to rescue the anti-proliferative effect of HDAC2 degradation **(Fig. 2j and Extended Data Fig. 2h)**, providing an orthogonal validation that *HDAC1* deletion is necessary to create this HDAC2 vulnerability.

To identify the direct transcriptional consequences of HDAC2 degradation, we performed thiol(SH)-linked alkylation for the metabolic sequencing of RNA (SLAM)-seq^41^, which quantifies changes in the abundance of newly synthesized mRNAs. Both HDAC2 degradation and HDAC1/2 inhibition (ACY-957^42^) predominantly upregulated transcriptional activity, producing tightly correlated SLAM-seq responses that emphasize the importance of HDAC2 function in an *HDAC1*-deficient background **(Fig. 3a,b)**. Interestingly, genes with HDAC2 bound at their promoter were not preferentially upregulated by HDAC2 degradation compared to those without HDAC2 **(Extended Data Fig. 3a)**. This prompted us to inspect promoter-distal sites of HDAC2 enrichment using ROSE2 (ranking of super enhancer 2), which was previously developed to identify super enhancers and their putative target genes^43^. From this analysis, we were able to classify genes as being proximal to sites of typical HDAC2 enrichment (94% of promoter-distal HDAC2-bound sites) or disproportionately high HDAC2 enrichment (6% of sites), which we termed asymmetrically bound sites **(Fig. 3c)**. After integrating these data with the 2-h SLAM-seq, we found that genes associated with typical or asymmetric sites were upregulated less than other genes in response to dTAG-13 treatment **(Fig. 3d)**. This effect, which did not occur following HDAC1/2 inhibition, was most pronounced at genes associated with asymmetric sites, which were nearly unchanged by dTAG-13 treatment **(Fig. 3d)**. Given the otherwise similar transcriptional response to HDAC1/2 inhibition and HDAC2 degradation, we were intrigued by this apparent difference at super enhancers.

**Figure 3:**
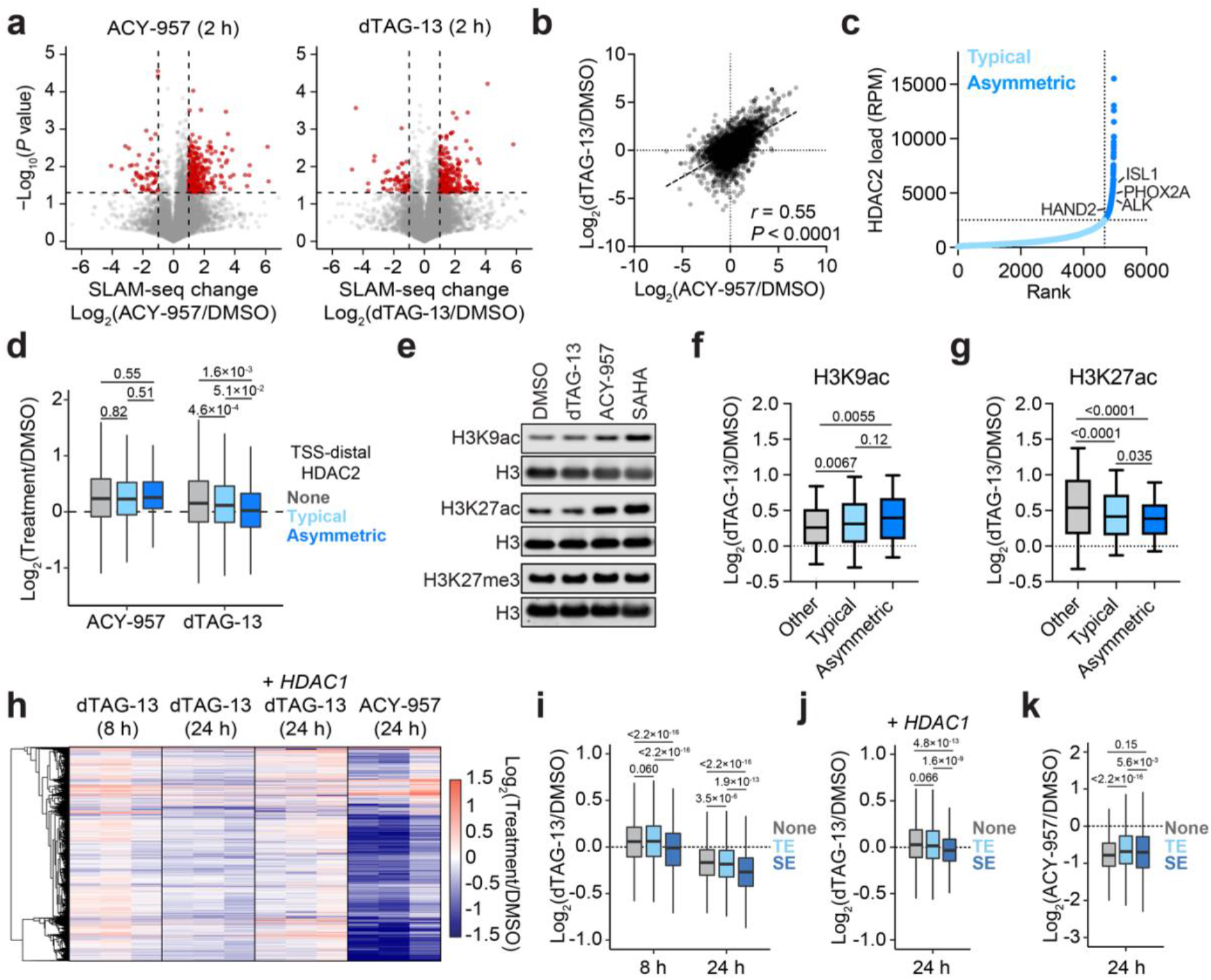
HDAC2 degradation dysregulates transcription. **a**, Volcano plots depicting changes in newly synthesized mRNAs (SLAM-seq) following 2-h treatments of 5 μM ACY-957 (left) or 500 nM dTAG-13 (right). *n* = 3. **b**, Correlation of SLAM-seq changes induced by ACY-957 and dTAG-13 treatments. *P* value was determined by Pearson correlation coefficient *(r). n* = 13,199. **c**, Asymmetric genomic localization of HDAC2 is depicted by plotting the rank-ordered CUT&RUN signal at each HDAC2-occupied site. **d**, Boxplot of changes in newly synthesized mRNAs (SLAM-seq). Genes are classified as not being associated with TSS-distal HDAC2 sites (white, n = 11,210), being associated with typical TSS-distal HDAC2 sites (light blue, n = 1,826), or being associated with asymmetric TSS-distal HDAC2 sites (dark blue, n = 163). **e**, Immunoblot detection of H3K9ac, H3K27ac, and H3K27me3 following 2-h treatments of DMSO, dTAG-13 (500 nM), ACY-957 (5 μM), or SAHA (5 μM). **f**, Boxplots of changes in H3K9ac CUT&RUN signal following 2-h dTAG-13 treatment (500 nM) at H3K9ac-enriched sites that overlap with TSS-distal HDAC2-typical sites (light blue, *n* = 882), with TSS-distal HDAC2-aysmmetric sites (dark blue, *n* = 133), and other sites (grey, *n* = 13,984). **g**, same as (**f**) but for H3K27ac (grey, *n*= 9,863; light blue, *n* = 2,149; dark blue, *n* = 280). Boxes represent 25-75 percentiles with whiskers extending to 10-90 percentiles. **h**, Heatmap of changes in total mRNA abundance (3’-end mRNA-seq) following dTAG-13 treatment (500 nM) or ACY-957 (5 μM) treatments in BE(2)-C-HDAC2-dTAG cells or BE(2)-C-HDAC2-dTAG overexpressing *HDAC1* cells with triplicates for each condition. **i**, Boxplots of changes in total mRNA abundance for genes not associated with an enhancer (grey, *n* = 7,588), associated with typical enhancers (light blue, *n* = 5,026), or associated with super enhancers (dark blue, *n* = 585). **j,k** As in (**i**) but with 24-h dTAG-13 treatment (500 nM) in BE(2)C-HDAC2-dTAG cells overexpressing *HDAC1* (**j**) or ACY-957 treatment (5 μM) in BE(2)C-HDAC2-dTAG cells (**k**). In volcano plots, *P* values were determined by two-tailed Student’s *t-*test. In boxplots, *P* values were determined by two-tailed Welch’s *t*-test. Analyses of gene expression changes were restricted to active genes (total CPM > 3, see **Methods**). Unless specified, boxplots represent 25-75 percentile with whiskers extending 1.5 interquartile range (IQR).

Although the hemizygous deletion of *HDAC1* makes it unable to compensate for HDAC2 degradation on a global scale **(Fig. 3a,b)**, we considered that residual HDAC1 activity might be able to contribute sufficient deacetylase function specifically at asymmetric sites. This would potentially explain why genes associated with asymmetric sites were not coordinately upregulated like we observed for other genes. To investigate this possibility, we measured the effect of dTAG-13 treatment on H3K9ac and H3K27ac levels, two well-established HDAC1/2 substrates^3,44^. While we observed minimal changes by immunoblot analysis **(Fig. 3e)**, we detected a slight increase of H3K9ac and H3K27ac abundance by CUT&RUN **(Extended Data Fig. 3b)**. The increase in H3K27ac was indeed muted at typical and asymmetric HDAC2 sites, but this was not the case for H3K9ac **(Fig. 3f,g)**. Since both modifications are known to be regulated by HDAC1/2, it is as-yet unclear if the difference in H3K27ac reflects residual HDAC1 activity or some other mechanism. Another possibility, not mutually exclusive with the first, is that the differences we observe with asymmetric sites might be the result of disrupting a non-enzymatic function of HDAC2 that is not addressed by HDAC1/2 inhibition. HDAC1 and HDAC2 are members of several multiprotein complexes, including the NuRD complex, which regulates chromatin accessibility through the nucleosome remodeler, CHD4. A recent study demonstrated that CHD4/NuRD-mediated chromatin remodeling positively regulates the transcription of genes associated with super enhancers^45^, which share considerable overlap with sites of asymmetric HDAC2 localization **(Extended Data Fig. 3c)**. Therefore, HDAC2 degradation might negatively impact NuRD function at super enhancers, which would, in principle, result in suppression of NuRD super enhancer target genes. This effect, combined with the more widespread transcriptional derepression caused by loss of HDAC catalytic activity, might also explain why most genes are upregulated by HDAC2 degradation, except those associated with asymmetric HDAC2 sites **(Fig. 3d)**.

To determine how super-enhancer-associated transcripts are regulated in response to HDAC2 degradation over time, we performed kinetic gene expression profiling experiments using 3’-end mRNA-seq to detect changes in total transcript abundance. After 8 hours of dTAG-13 treatment, we observed a general increase of transcriptional activity **(Fig. 3h and Extended Data Fig. 3d)**, which is consistent with the general upregulation we observed by SLAM-seq after 2 hours **(Fig. 3a)**. Indeed, the genes that were significantly upregulated after 2 hours were preferentially upregulated after 8 hours **(Extended Data Fig. 3g)**. However, after 24 and 72 hours, transcriptional activity was generally suppressed **(Fig. 3h and Extended Data Fig. 3e,f)**, consistent with previous reports that class I HDAC inhibition causes only transient transcriptional upregulation^46^. Again, we saw no obvious difference in the expression of genes based on whether HDAC2 is bound at the promoter **(Extended Data Fig. 3h)**, but super-enhancer-associated transcripts were distinctly affected at every time point **(Fig. 3i and Extended Data Fig. 3l)**. That is, despite otherwise widespread increases in transcriptional activity after 8 hours, the transcription of SE-linked genes was relatively unchanged. Then, after 24 and 72 hours, when transcriptional downregulation becomes predominant, SE-linked genes were more severely suppressed than others **(Fig. 3i and Extended Data Fig. 3l)**. Altogether, these data suggest that SE-associated transcripts, which are enriched with lineage-specific dependencies **(Extended Data Fig. 3m)**, are potently inhibited by HDAC2 degradation **(Fig. 3i and Extended Data Fig. 3n)**. Importantly, exogenous expression of *HDAC1* rescued the transcriptional effects caused by HDAC2 degradation **(Fig. 3j and Extended Data Fig. 3i)**. After 24 hours of dTAG-13 treatment in cells overexpressing BE(2)-C-HDAC2-dTAG cells overexpressing *HDAC1,* we did not detect gross changes in transcriptional activity and the preferential repression of super enhancer-associated transcripts was substantially muted **(Fig. 3j and Extended Data Fig. 3i)**. We next performed 3’-end mRNA-seq following a 24-h treatment of ACY-957 to determine if the preferential suppression of SE-linked genes is specific to degradation versus inhibition **(Fig. 3k and Extended Data Fig. 3j)**. In contrast to the correlated effects of HDAC1/2 inhibition and HDAC2 degradation at 2 hours by SLAM-seq **(Fig. 3b)**, changes in total mRNA abundance after 24 hours were poorly correlated **(Extended Data Fig. 3k)** and ACY-957 did not preferentially downregulate SE-linked genes like we observe following HDAC2 degradation **(Fig. 3j)**.

Returning to the idea that HDAC2 degradation may impact NuRD function at super enhancers, we wondered whether loss of HDAC2 may destabilize the complex and lead to its degradation. This phenomenon, whereby chemically induced degradation results in destabilization of other subunits within a large multiprotein complex, has been reported for PRC2, BAF (by both direct-acting PROTACs and dTAG-based degradation), and some HDAC-associated complexes^47–50^. The NuRD complex is likely susceptible to this effect as well. Although the mechanism is as-yet unclear, a recent study reported the discovery of an electrophilic small molecule that causes the coordinated degradation of several NuRD subunits in T cells^51^. Based on these examples, we probed the effect of HDAC2 degradation on protein stability using unbiased quantitative expression proteomics **(Fig. 4a,b)**. As expected, loss of HDAC2 abundance is the most profound change that we observed in this dataset, confirming the immediate and selective effects of the dTAG system in BE(2)-C-HDAC2-dTAG cells **(Fig. 4a)**. Interestingly, gene set enrichment analysis (GSEA) revealed that other HDAC-associated proteins are also preferentially lost after 2- and 24-h dTAG-13 treatments **(Fig. 4c,d and Extended Data Fig. 4a** Reactome HDACs deacetylate histones).

**Figure 4:**
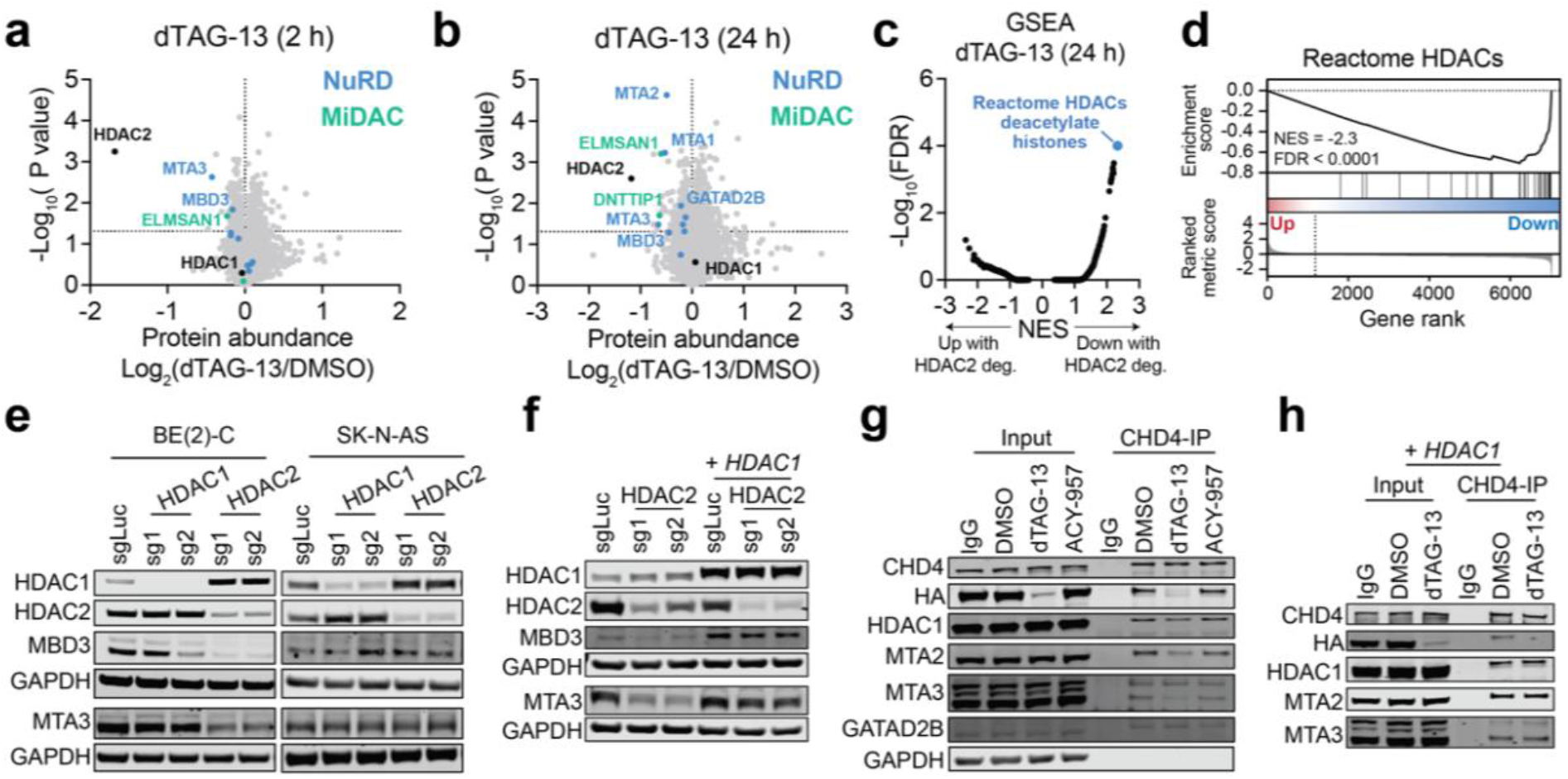
HDAC2 degradation destabilizes the NuRD complex in HDAC1-deficient neuroblastoma cells. **a,b** Volcano plots of changes in protein abundance determined by unbiased quantitative proteomics following dTAG-13 (500 nM) treatment for 2 h (**a**) or 24 h (**b**). DMSO *n* = 2, dTAG-13 *n* = 3. *P* values were determined by two-tailed Student’s *t*-test. **c**, Gene set enrichment analysis (GSEA) for protein abundance changes after 24-h dTAG-13 treatment. NES, normalized enrichment score. **d**, GSEA of Reactome HDACs gene set for protein abundance changes after 24-h dTAG-13 treatment. **e**, Protein levels of NuRD subunits following genetic disruption of *HDAC1* and *HDAC2* by CRISPR/Cas9 in BE(2)-C (left) and SK-N-AS (right) determined by immunoblot analysis. **f,** As in (**d**) but in BE(2)-C cells with or without *HDAC1* overexpression. **g**, Co-immunoprecipitation of CHD4 in BE(2)-C-HDAC2-dTAG cells with 2-h dTAG-13 (500 nM) and ACY-957 (5 μM) treatments. **h**, As in (**f**) but with 2-h dTAG-13 treatment in BE(2)-C-HDAC2-dTAG cells with *HDAC1* overexpression.

Closer inspection of these data revealed that several members of two HDAC1/2-containing complexes, mitotic deacetylase complex (MiDAC) and NuRD, are depleted by dTAG-13 treatment **(Fig. 4a,b)**. This occurs without any change in gene expression, confirming it is a post-transcriptional effect **(Extended Data Fig. 4b)**. Repeating these experiments using immunoblot detection, we were able to reproduce the depletion of NuRD subunits but failed to identify changes to the MiDAC complex **(Extended Data Fig. 4c,d)**. We therefore focused our efforts on studying NuRD structure and function in response to HDAC2 degradation. The NuRD complex consists of subunits that catalyze ATP-dependent chromatin remodeling (CHD3/4) and histone deacetylation (HDAC1/2), accompanied by several scaffolding and chromatin-binding proteins, including MTA1/2/3, GATAD2A/B, MBD2/3 and RBBP4/7^52^. MTA paralogs bind directly to HDAC1 and HDAC2, forming a subcomplex together with RBBP4/7, while CHD3/4 and GATAD2A/B form the other half of the complex^53,54^. MBD2/3 are thought to bridge the two halves together to constitute a complete structure and are thus critical for complex integrity^53,54^. Moreover, as several paralogous members of the complex are mutually exclusive, distinct complex compositions can preferentially affect certain cell types or cellular processes^55^. Mechanistically, NuRD degradation might be caused by direct PROTAC-facilitated ubiquitination of non-HDAC subunits or occur as a secondary consequence of protein complex destabilization following HDAC2 degradation. We strongly favor the latter explanation, as we detect decreased abundance of NuRD subunits following CRISPR/Cas9-mediated disruption of *HDAC2* **(Fig. 4e)**, and because more subunits were affected after 24 hours of dTAG-13 treatment than after 2 hours **(Fig. 4b)**.

We next asked whether NuRD degradation was specific to cells with compromised levels of HDAC1. Indeed, CRISPR/Cas9-mediated disruption of *HDAC2* in the *HDAC1*-diploid SK-N-AS cells did not affect the abundance of other NuRD subunits **(Fig. 4e)** and the effect in BE(2)-C cells could be rescued by overexpression of *HDAC1* **(Fig. 4f)**. We therefore hypothesized that HDAC2 degradation might result in compromised intra-complex interactions in cells without sufficient levels of HDAC1 to stabilize the complex. To probe NuRD interactions, we performed CHD4 co-immunoprecipitations following a 2-h dTAG-13 treatment and noted weakened interactions between CHD4 and HDAC1, MTA2 and MTA3, but not between CHD4 and GATAD2B **(Fig. 4g)**. These data indicate that the HDAC/MTA subcomplex is immediately dissociated from the CHD4 subcomplex upon HDAC2 degradation, which results in degradation of MTA and MBD subunits. Moreover, the impact of HDAC2 degradation on NuRD interactions was rescued by overexpression of *HDAC1* **(Fig. 4h)**, confirming that the effect is dependent on diminished HDAC1 protein abundance.

To determine if the effect of HDAC2 degradation on NuRD stability impacts its chromatin remodeling activity, we first mapped the genomic localization of MBD3 and CHD4 in BE(2)-C cells by CUT&RUN **(Fig. 5a)**. In comparison to HDAC2, which occupies both promoters and enhancers, MBD3 and CHD4 show a predominant localization to enhancers **(Fig. 5a and Extended Data Fig. 5a)**. About 60% of HDAC2-bound sites overlap with either MBD3 or CHD4 **(Fig. 5b)** and these sites (HDAC2-NuRD) are preferentially associated with enhancers **(Fig. 5c)**. In fact, approximately 70% of super enhancers and 16% of typical enhancers are associated with HDAC2-NuRD **(Fig. 5d)**. In contrast, HDAC2-bound sites that do not co-localize with MDB3/CHD4 consist mostly of promoter regions **(Fig. 5c)**. In general, we interpret this to indicate that HDAC2 localization to enhancers is commonly associated with the NuRD complex, whereas its localization to promoters is associated with other complexes. For example, at *ISL1,* which is a highly selective neuroblastoma dependency that is downregulated by HDAC2 degradation **(Extended Data Fig. 1a and Extended Data Fig. 3n)**, we observe HDAC2 bound at the promoter, but no binding of MBD3 or CHD4 **(Fig. 5e)**. In contrast, all three members of the NuRD complex are highly enriched at its proximal super enhancer **(Fig. 5e)**.

**Figure 5:**
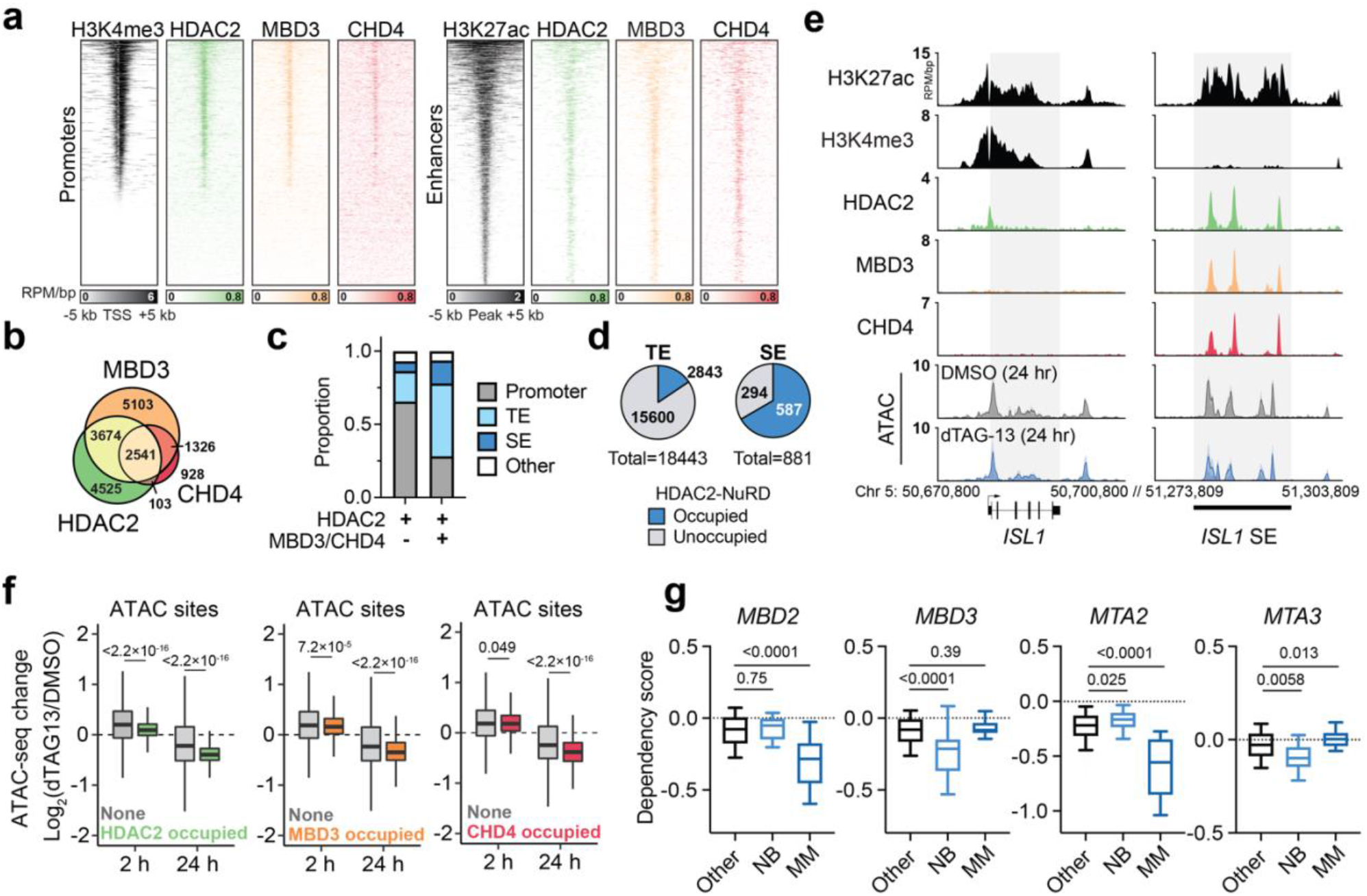
HDAC2 degradation compromises NuRD function and exploits cancer-specific NuRD dependencies. **a**, Heatmaps of CUT&RUN signals for HDAC2, MBD3, and CHD4 mapped to promoters (left, ranked by H3K4me3 ChIP signal from GSM2113519) and enhancers (right, ranked by H3K27ac ChIP signal from GSM2113518). **b,** Overlap of HDAC2-, MBD3-, and CHD4-bound sites. HDAC2 bound-sites shared with MBD3 and/or CHD4 are defined as HDAC2-NuRD sites. **c**, HDAC2-only sites (HDAC2 sites that do not overlap MBD3 or CHD4) mainly localize to promoters while HDAC2-NuRD predominantly occupy enhancers. **d**, Proportion of enhancers and super enhancers that are occupied by HDAC2-NuRD. **e**, Gene track representation of ChIP-seq, CUT&RUN, or ATAC-seq signal at the *ISL1* locus (left) and its proximal enhancer (right). ATAC-seq signal is shown as meta-track representation of triplicates following 24-h DMSO control (grey) or 500 nM dTAG-13 treatment (blue). **f**, Boxplots of chromatin accessibility changes grouped by HDAC2 occupancy (left) (2 h, unoccupied *n* = 97,849, occupied *n* = 9,581; 24 h, unoccupied *n* = 94,967, occupied *n* = 9,573), MBD3 occupancy (middle) (2 h, unoccupied *n* = 96,391, occupied *n* = 11,039; 24 h, unoccupied *n* = 93,608, occupied *n* = 10,932), or CHD4 occupancy (right) (2 h, unoccupied *n* = 103,049, occupied *n* = 4,381; 24 h, unoccupied *n* = 100,223, occupied *n* = 4,317) at ATAC-seq sites. Boxes represent 25-75 percentiles with whiskers extending 1.5 IQR. *P* values were determined by two-tailed Welch’s *t-*test. **g**, Boxplots of *MBD2, MBD3, MTA2,* and *MTA3* dependency scores in neuroblastoma (*n* = 34), multiple myeloma (*n* = 21), or other cell lines (*n* = 999). Boxes represent 25-75 percentiles with whiskers extending to 10-90 percentiles. *P* values were determined by two-tailed Student’s *t*-test.

Next, we performed assay for transposase-accessible chromatin (ATAC)-seq^56^ following 2-h and 24-h exposures to dTAG-13 to determine if NuRD chromatin remodeler activity is impacted by HDAC2 degradation **(Extended Data Fig. 5b)**. Similar to the transcriptional effects we observed after 2 hours, acute HDAC2 degradation causes widespread increases of chromatin accessibility **(Extended Data Fig. 5b)**. However, HDAC2-bound sites of the genome are significantly less affected by this increase **(Fig. 5f)**, which we noted was similar to the observation that genes associated with asymmetric HDAC2 sites were less upregulated at 2 hours **(Fig. 3d)**. After 24 hours, chromatin accessibility was widely diminished, with pronounced effects evident at HDAC2-bound sites **(Fig. 5f and Extended Data Fig. 5b)**. This decrease in chromatin accessibility that follows extended HDAC2 degradation is likewise consistent with the transcriptional suppression we observe at 24 hours **(Extended Data Fig. 3e)**, but contrary to past reports that link HDAC inhibition (both Class I and pan-HDAC) with increased chromatin accessibility^57,58^. Therefore, we wondered whether the differential regulation of accessibility at HDAC2-bound sites reflects the opposing effects of HDAC2 catalytic activity, which represses chromatin accessibility, and NuRD remodeler function, which maintains open chromatin structures at super enhancers^45^. Indeed, we find that accessibility at MBD3- and CHD4-bound sites become preferential repressed following HDAC2 degradation for 24 hours **(Fig. 5f)**. These data indicate that HDAC2 degradation not only disrupts NuRD integrity, but also disrupts its chromatin remodeler function, which is altogether consistent with the pronounced suppression of SE-associated transcripts we detect **(Fig. 3i and Extended Data Fig. 3l)**.

Interested in how NuRD degradation might be expected to impact cancer cell survival, we analyzed the dependency map and noted that specific paralogs within the NuRD complex, but not other HDAC1/2-associated complexes, register as highly selective dependencies in neuroblastoma and multiple myeloma on DepMap **(Fig. 5g and Extended Data Fig. 5c,d)**. Interestingly, discrete sets of paralogs are selectively required by neuroblastoma and others are required by multiple myeloma. For example, *MBD2* and *MTA2* are selectively required for multiple myeloma, whereas *MBD3* and *MTA3* are selectively required for neuroblastoma **(Fig. 5g)**. Using DepMap to identify co-dependencies of these NuRD subunits, we find that *MBD2* and *MTA2* are most highly correlated with each other, followed by several top multiple myeloma dependencies, like *IRF4* **(Extended Data Fig. 5e)**. On the other hand, *MBD3* and *MTA3* dependency scores are highly correlated with HDAC2 and other selective neuroblastoma dependencies, such as *MYCN* and *ISL1* **(Extended Data Fig. 5e)**. These correlations suggest the possibility that functionally distinct NuRD subcomplexes are relied on in the two cell types. This would be consistent with the known capacity of the NuRD complex to form functionally non-redundant subcomplexes through differential assembly of paralogous subunits^55^. We used unsupervised clustering of all NuRD dependency scores across all cancer cell lines on DepMap to identify functional trends among NuRD paralogs, as this type of co-dependency analysis has previously been used to identify functionally and physically distinct protein complex assemblies^59,60^. This revealed a cluster containing *MBD2, MTA2,* and *HDAC1,* and another containing *MBD3, MTA3,* and *HDAC2* **(Extended Data Fig. 5f)**, which tracks with the former being required by multiple myeloma and the latter being required by neuroblastoma. It is therefore attractive to speculate that these cancer types rely on distinct NuRD subcomplex compositions for tumor maintenance. Since these NuRD dependencies are degraded following loss of HDAC2, we conclude that exploiting HDAC1/2 synthetic lethality represents an effective strategy to target lineage-specific NuRD vulnerabilities.

## Discussion

This study applied a generalizable framework for identifying synthetic lethal genetic interactions using genetic dependency maps made publicly available pre-publication. We identified *HDAC1/2* as a reciprocal synthetic lethality caused by common chromosomal deletions observed in multiple human cancers. Using genetic perturbations, chemically responsive degradation, and genetic rescue experiments, we demonstrate that hemizygous deletions of *HDAC1* or *HDAC2* are necessary and sufficient to sensitize cells to loss of the unaffected paralog. The dTAG system for chemically induced degradation was particularly useful for determining the mechanisms underlying this synthetic lethality, ultimately revealing an effect on NuRD complex stability, chromatin accessibility, and cell-type-specific transcriptional regulation.

Collectively, we demonstrate that targeting *HDAC1/2* synthetic lethality compromises the structure and function of the NuRD complex, leading to the degradation of NuRD subunits that are selectively essential in neuroblastoma. Our study follows the discovery of a conceptually similar synthetic lethality between *NXT1* and *NXT2* in neuroblastoma and rhabdomyosarcoma^19^. When NXT1 is lost from tumors with compromised *NXT2* expression, the NXT1/2 binding partner, NXF1, which is an essential protein, is subjected to degradation^19^. In our study, we find that loss of HDAC2 in cells with hemizygous deletion of *HDAC1* results in destabilization of the NuRD complex and degradation of several neuroblastoma-specific dependencies contained therein. Additionally, CHD4, which is pan-essential, can no longer interact with several NuRD subunits, likely indicating that it is functionally compromised in these cells. This inference is supported by the loss of chromatin accessibility that we observe at CHD4-bound sites after 24 hours of HDAC2 degradation, as previous CHD4 loss-of-function genetic experiments revealed CHD4 as a positive regulator of chromatin accessibility^45^. Therefore, we conclude that perturbing the stability and/or function of essential proteins (either lineage-specific or pan-essential) by exploiting synthetic lethality between paralogs is likely to be a generalizable framework for discovering new anti-cancer drug targets.

The fast kinetics of the dTAG system allowed us to dissect HDAC2 function with exceptional temporal resolution, revealing the effects of HDAC2 degradation at early and late time points. Immediately following the loss of HDAC2, we observed transient transcriptional upregulation and increased chromatin accessibility. This gave way to transcriptional repression, coinciding with NuRD destabilization and chromatin compaction at NuRD-binding sites. Transcriptionally, the preferential downregulation of SE-linked genes, which are regulated by CHD4/NuRD^45^, results in the repression of many genes that are selectively required for neuroblastoma survival. These findings join an increasing body of evidence that links HDAC/NuRD to transcriptional activation, which contrasts with the paradigmatic role of HDACs as transcriptional repressors^61,62^.

Our findings provide further motivation for the ongoing effort to develop direct-acting HDAC1/2 PROTACs^47,63^. To effectively capitalize on this synthetic lethality therapeutically, it will be necessary to develop selective degraders that only target one paralog or the other, thereby avoiding the toxicity associated with combined HDAC1/2 loss of function. Fortunately, it has been well demonstrated that PROTACs can achieve greater selectivity than their parental compounds, with multiple examples now of degraders that differentiate between two highly similar paralogs^64^. It may therefore be possible to develop paralog-selective degraders from ligands that bind both HDAC1 and HDAC2. While these proteins have proven difficult targets for PROTAC development^47^ in general, we hope that these data will inspire renewed motivation to develop effective PROTACs and/or molecular glue degraders.

## Materials and Methods

### Cell culture and lentivirus production

BE(2)-C (DMEM supplemented with 10% fetal bovine serum (FBS) and Gibco Antibiotic-Antimycotic), SK-N-AS (DMEM supplemented with 10% FBS and Gibco Antibiotic-Antimycotic), and OCI-AML2 (RPMI supplemented with 10% FBS and Gibco Antibiotic-Antimycotic) cell lines were provided by the laboratory of Prof. James E. Bradner. MM.1S (RPMI supplemented with 10% FBS and Gibco Antibiotic-Antimycotic) cells was provided by the laboratory of Prof. Christopher Ott. RPMI-8226 (RPMI supplemented with 15% FBS and Gibco Antibiotic-Antimycotic) cells was purchased from DSMZ. Lenti-X 293T (DMEM supplemented with 10% FBS and Gibco Antibiotic-Antimycotic) cells were purchased from Takara for lentivirus production. All cell lines were tested negative for mycoplasma infections regularly. Lentiviral packaging plasmids pMD2.G (a gift from Didier Trono, Addgene plasmid #12259; http://n2t.net/addgene:12259; RRID:Addgene_12259), psPAX2 (a gift from Didier Trono, Addgene plasmid #12260; http://n2t.net/addgene:12260; RRID:Addgene_12260), and the lentiviral expression plasmid were co-transfected to Lenti-X 293T cells to produce corresponding lentivirus. Supernatants with viral particles were harvested at 48 and 72 hours after transfection, filtered with 0.22 μm membrane and concentrated by 50-fold with Lenti-X Concentrator (Takara, #631232). All cells were transduced by spinoculation at 2000 rpm for 1 hour at room temperature supplemented with 8 μg/mL polybrene. For MM.1S and RPMI-8226 cells polybrene was removed 8 hours after the spinoculation to prevent toxicity.

### Analyses of datasets from DepMap

Unless specified, all data were obtained from 21Q4 datasets release including CRISPR_genetic_effect (Chronos), CCLE_expression, and CCLE_gene_cn. For HDAC1/2 copy number analyses in **Fig. 1h&l** we used Log2(relative to ploidy + 1) of 0.7 as cutoff for hemizygous deletion to include all cell lines potentially harboring deleted HDAC1/2, based on the calculation that 1 copy of the gene equals Log2(relative to ploidy + 1) of 0.58.

### Analyses of datasets from cBioportal^34^

The analysis of neuroblastoma patient data is based upon data generated by the Therapeutically Applicable Research to Generate Effective Treatments (TARGET) initiative, phs000218, managed by the NCI. Specifically, data used in this study were obtained from Pediatric Neuroblastoma (TARGET, 2018) (cancer study identifier: nbl_target_2018_pub). Information about TARGET can be found at http://ocg.cancer.gov/programs/target. DLBCL patient data were obtained from Lymphoid Neoplasm Diffuse Large B-cell Lymphoma (TCGA, Firehose Legacy) (cancer study identifier: dlbc_tcga). Source data is from http://gdac.broadinstitute.org/runs/stddata_2016_01_28/data/DLBC/20160128/.

### Immunoblotting

Cells were resuspended with CelLytic M buffer (Sigma-Aldrich, #C2978) supplemented with 1X Halt Protease Inhibitor Cocktail (Thermo Scientific, #78430) and 0.1% benzonase endonuclease (Millipore, #70746) for whole cell lysates. Lysates were incubated on ice for 30 mins with occasional vortexing and cleared by 13,000 *g* centrifugation at 4° C for 10 mins. The total protein concentration was then measured with Pierce BCA protein assay kit (Thermo Scientific, #23225). Samples on one blot were normalized to the same total protein content. To extract histones cells were resuspended with 0.5% Triton X 100 in PBS supplemented with 1X Halt Protease Inhibitor Cocktail (Thermo Scientific, #78430) and incubated on ice for 10 mins. The pellets were collected after 2,000 rpm centrifugation at 4° C for 10 mins. Histones were extracted with 0.2 N HCl at 4° C overnight. The protein content was then determined by Bradford assay (Thermo Scientific, #23200) and equal amount of protein across samples was run on the same blot. Protein bands were detected by fluorescently labeled infrared secondary antibodies (IRDye) on the Odyssey CLx Imager (LI-COR). The antibodies used are all commercially available: anti-CHD4 (Cell Signaling Technology, #12011S), anti-HDAC1 (Cell Signaling Technology, #34589S), anti-HDAC2 (Cell Signaling Technology, #5113S), anti-HA (Cell Signaling Technology, #3724S), anti-MBD3 (Cell Signaling Technology, #99169S), anti-MTA2 (Bethyl, #A300-395A), anti-MTA3 (Proteintech, #14682-1-AP), anti-H3 (Cell Signaling Technology, #14269S), anti-H3K9ac (Cell Signaling Technology, #9649S), anti-H3K27ac (Abcam, #ab4729), anti-H3K27me3 (Cell Signaling Technology, #9733S).

### Engineering of dTAG cell line

The HDAC2-dTAG cell line was engineered using the PITCh system^65^, as previously described^39^, using microhomology-mediate end joining (sgRNA: 5’-GTTGCTGAGCTGTTCTGATT-3’) to insert a linker-FKBP12^F36V^-2xHA-P2A-BSD^R^ cassette into the C-terminus of *HDAC2*. BE(2)-C cells were transfected with both CRIS-PITCh plasmids, selected by 10 μg/mL Blasticidin S HCl, and clonally expanded to confirm biallelic knock-in by PCR and immunoblot.

### Cellular proliferation

BE(2)-C cells were plated into 12-well plates with 12,000 cells in 1 ml medium per well. DMSO and compounds were added 1:1000 in triplicates. Every 3 to 4 days cells were trypsinized, counted by Countess automated cell counter (Invitrogen), and re-plated at the concentration of 12,000 cells/ml with fresh DMSO or compounds.

### CRISPR-Cas9 competitive growth assays

Cas9 expressing cells were created by transduction of lentiCas9-Blast (a gift from Feng Zhang, Addgene plasmid #52962; http://n2t.net/addgene:52962; RRID:Addgene_52962) virus and selected by 10 μg/mL Blasticidin S HCl. sgRNA sequences were cloned into the LRG backbone^66^ (a gift from Christopher Vakoc, Addgene plasmid #65656; http://n2t.net/addgene:65656; RRID:Addgene_65656) for co-expression of an sgRNA and GFP. A modified version of the LRG vector with mCherry replacing GFP was also developed for this study. Cas9-expressing cells were transduced in triplicate with the corresponding sgRNA-LRG virus at 30-60% efficiency in 96-well plates. For two-color experiments, cells were transduced with GFP-sgRNA1 and mCherry-sgRNA2 simultaneously to achieve ~25% of each population (non-fluorescent, GFP or mCherry positive, GFP and mCherry positive). Transduced cells were passaged and subjected to flow cytometry to measure GFP- and/or mCherry-positive percentage every 3 to 4 days. sgRNA sequences, which were taken from the Brunello sgRNA library^67^, are as follows: AAVS1 5’-GGGGCCACTAGGGACAGGAT-3’; RPS19 5’-GTAGAACCAGTTCTCATCGT-3’; Luciferase 5’-CCCGGCGCCATTCTATCCGC-3’; HDAC1-sg1 5’-CATCCGTCCAGATAACATGT-3’; HDAC1-sg2 5’-TGAGTCATGCGGATTCGGTG-3’; HDAC1-sg3 5’-GCACCGGGCAACGTTACGAA-3’; HDAC1-sg4 5’-GGAGATGTTCCAGCCTAGTG-3’; HDAC2-sg1 5’-GATGTATCAACCTAGTGCTG-3’; HDAC2-sg2 5’-TACAACAGATCGTGTAATGA-3’;HDAC2-sg3 5’-CCTCCTCCAAGCATCAGTAA-3’; HDAC2-sg4 5’-TCAAAGAGTCCATCAAACAC-3’. For sgAAVS1, cutting efficiency was determined by quantitative sequence trace decomposition (tracking of indels by decomposition software; TIDE) ^32^ following PCR amplification and Sanger sequencing of the cut site (5’-AGGTGGGGGTTAGACCCAAT-3’, 5’-CTTCTCCGACGGATGTCTCC-3’). The efficiency of all other sgRNAs was confirmed by immunoblot.

### Generation of knockout populations

To create loss-of-function knockout populations, Cas9-expressing cells were transduced with the corresponding sgRNA-LRG vector to achieve 90-100% efficiency, as confirmed by flow cytometry. Population-level knockouts were validated by immunoblot of the corresponding protein.

### Immunoprecipitation

50 million cells were collected after drug treatment and washed twice with 5 mL cold PBS. Cells were then lysed in 1 ml lysis buffer (50 mM HEPES, 150 mM NaCl, 1.5 mM MgCl2, 1% NP40, 125 U benzonase, and 1X Halt Protease Inhibitor Cocktail) with mild sonication and incubated on a rotating platform at 4° C for 3 hours. The supernatant was collected following 18,000 *g* centrifugation at 4° C for 20 mins. The protein content for each sample was quantified by BCA assay and normalized to the same level. For CHD4 immunoprecipitations, 50 μL of Dynabeads protein G (Invitrogen, #10003D) were combined with 10 mg antibody (anti-CHD4, Abcam, #ab70469-10) and rotated at room temperature for 30 mins to make antibody-beads conjugates, which were then washed three times supplemented with 1 mL PBS with 0.05% Tween20. For HA immunoprecipitations, 30 μL Pierce anti-HA Magnetic Beads (Thermo Fisher Scientific, #88836) were used for each sample. The antibody-bead conjugates were added to the protein lysate and rotated at 4° C overnight for immunoprecipitation. The beads were washed with 1 mL cold wash buffer (50 mM HEPES, 150 mM NaCl, and 0.2% NP40) for three times and eluted with 1X Bolt LDS Sample Buffer (Invitrogen, #B0007) supplemented with 2.5% 2-Mercaptoethanol by incubation at 95° C for 10 mins. Eluted samples were subjected to immunoblot.

### Animal experiments

Mouse experiments were performed in accordance with the Institutional Animal Care and Use Committee and conform to the guidelines for the care and use of laboratory animals approved at The Scripps Research Institute (protocol #18-031-01). Xenografts were established in six-week-old female NOG (NOD.Cg-*Prkdc^scid^ Il2rg^tm1Sug^/JicTac)* mice (Taconic) by flank injection of 0.8 million cells in 50% Matrigel (BD Biosciences) in DMEM (Gibco, ThermoFisher Scientific), total volume 50 μL. Experimental groups consisted of 5 mice, with two flank tumors each. Tumor burden was assessed by measuring tumor length and width using digital calipers once per week for the first two weeks then three times per week for up to six weeks. Caliper measurements of the longest axis were used to calculate the diameter. Serial caliper measurements of perpendicular axes were used to calculate tumor volume by the following formula: (short diameter × 2) × (long diameter) × 0.5. Mice were sacrificed once the longest diameter of at least one flank tumor reached 15 mm. At the endpoint, tumors were collected for molecular studies.

### Cleavage Under Targets & Release Using Nuclease (CUT&RUN)

All CUT&RUN experiments were performed as previously described^40^ with minor modifications. Briefly, 50,000 cells were collected and washed twice with 1 mL wash buffer (20 mM HEPES-KOH, 150 mM NaCl, 0.5 mM spermidine, 1X HALT protease inhibitor cocktail) at room temperature for each sample. Concanavalin A coated magnetic beads (BioMag, #BP531, 40 μL per sample) were activated and used to capture cells by incubating at room temperature on a rotating platform for 15 mins. Cells were permeabilized on beads with 250 μL antibody buffer (20 mM HEPES-KOH, 150 mM NaCl, 0.5 mM spermidine, 0.02% digitonin, 2 mM EDTA, 1 mM EGTA, 1X HALT protease inhibitor cocktail) with 1:100 corresponding antibody (anti-CHD4, Cell Signaling Technology, #12011S; anti-HDAC2, Active motif, #31505; anti-HA, Cell Signaling Technology, #3724S; anti-MBD3, Cell Signaling Technology, #99169S; anti-H3K9ac, Cell Signaling Technology, #9649S; anti-H3K27ac, Abcam, #ab4729; anti-IgG, CiteAb, ABIN101961). Samples were incubated at 4° C for 2 hours on a rotating platform, washed with 1 mL cold digitonin buffer (20 mM HEPES-KOH, 150 mM NaCl, 0.5 mM spermidine, 0.02% digitonin) twice, and then incubated with 350 ng of pA-MNase (a gift from the Henikoff lab) in 250 μL of digitonin buffer at 4° C for 1 hour. Samples were then washed three times with 1 mL cold digitonin buffer, resuspended in 150 μl digitonin buffer, and placed on a metal block equilibrated to 0° C. The MNase digestion was activated with 2 mM CaCl2 and performed for 30 mins at 0° C with occasional mixing of the beads. The reaction was stopped by addition of 150 μL 2X stop buffer (340 mM NaCl, 20 mM EDTA, 4 mM EGTA, 0.02% digitonin, 0.05 mg/mL RNase A, 0.05 mg/mL glycogen). The CUT&RUN fragments were released from cells by incubation at 37° C for 30 mins and collected from the supernatant. Additional 2 μL of RNase A (10 mg/mL) was added to each sample and incubated at 37° C for 30 mins. 0.1% SDS and 0.2 mg/mL proteinase K were added for protein digestion performed at 70° C for 30 mins. The DNA fragments were collected by phenol-chloroform extraction and resuspended in 25 μL of DNase-free water. The fragment distribution was then checked by TapeStation (Agilent). Sequencing libraries were prepared using Thruplex DNA-seq kit (Takara, # R400675) and 10 μL of input material. Libraries were amplified with modified conditions to allow preferential amplification of the short target fragments from CUT&RUN (16 cycles of 98° C for 15 s and 60° C for 10 s). Libraries were then purified with AMPure XP beads (Beckman Coulter, #A63880) and sequenced on Illumina NextSeq 2000 (paired-end, 35-bp reads). The sequencing reads were aligned to human genome build hg19 and RefSeq genes by Bowtie2 with the following parameters: --local -very-sensitive-local --no-unal --no-mixed --no-discordant --phred33 -I 10 -X 700 --threads 12 -x. Using the IgG sample as background, the signal-enriched regions were identified by Model-based Analysis of ChIP-seq (MACS12) peak-finding algorithm (v1.4.1) with the *P* value threshold of 1e-9. For differential analyses across samples, the read density calculator Bamliquidator (v1.0)(http://github.com/BradnerLab/pipeline/wiki/bamliquidator) was used to map sequencing reads to designated loci with a 200 bp extension window in either direction. The read density per base pair was calculated with the normalization to the total number of million mapped read (RPM/bp). ROSE2 was used with default parameters to identify asymmetric binding by HDAC2 (https://github.com/linlabbcm/rose2). H3K27ac ChIP-seq in wild-type BE(2)-C (GEO ID: GSM2113518) and its input sample (GEO ID: GSM2113520) were obtained from a previous study^29^ and used to predict enhancers and super enhancers with ROSE2. H3K4me3 ChIP-seq in wild-type BE(2)-C (GEO ID: GSM2113519) from a previous study^29^ was used as a mark for active promoters.

### Thiol(SH)-linked alkylation for the metabolic sequencing of RNA (SLAM-seq)

SLAM-seq was performed as previously described^41^ using the SLAMseq Kinetics Kit (Lexogen, #061.24). Briefly, 1 million cells were pretreated with 0.1% DMSO, 500 nM dTAG-13, or 5 μM ACY-957 for 1 hour in triplicate. Newly synthesized mRNA was then labeled by addition of 100 μM 4-thiouracil (4sU) and cells were incubated for another hour. Total RNA for each sample was extracted with RNeasy miniprep kit (Qiagen) and quantified by Qubit RNA XR Assay kit (Invitrogen, # Q33223). 5 μg RNA was subjected to alkylation by 100 mM iodoacetamide. The alkylated RNA was purified by ethanol precipitation. 1 μg of RNA for each sample was used to prepare the library for sequencing with QuantSeq 3’ mRNA-Seq Library Prep Kit FWD for Illumina (Lexogen, #015) and sequenced on Illumina NovaSeq 6000 (single-end, 101-bp reads). The sequencing data was analyzed by SlamDunk^68^ with default settings to align to human genome build hg19, and to calculate counts per million (CPM) and T-to-C conversion rate at 3’ UTRs (built from ncbiRefSeqCurated table for hg19 accessible on UCSC table browser, including 29,870 3’ UTRs for 22,514 genes). Conversion rate of each transcript in each sample can be found in **Supplementary Table S1**.

### 3’ mRNA-seq with ERCC RNA Spike-In Mix normalization

Cells were plated in 6-well plates with equal number and treated in triplicates. Total RNA was extracted with RNeasy miniprep kit (Qiagen). 350 μL of lysis buffer (RLT) supplemented with 2.1 ng/mL of ERCC RNA Spike-In Control SIRV Set 3 (Lexogen, #051.0) for spike-in normalization was used to lyse cells for each sample. 1 μg of RNA for each sample was used to prepare sequencing library with QuantSeq 3’ mRNA-Seq Library Prep Kit FWD for Illumina (Lexogen, #015). The libraries of 8-h and 24-h DMSO and 500 nM dTAG-13 treatments were sequenced on Illumina NovaSeq (single-end, 101-bp reads); The libraries of 24-h DMSO and 5 μM ACY-957 treatments were sequenced on Illumina Nextseq 500 (single-end, 100-bp reads); The libraries of 72-h DMSO and 500 nM dTAG-13 treatments were sequenced on Illumina Nextseq 2000 (single-end, 75-bp reads). SlamDunk was used to align the sequencing reads to human genome build hg19 and ERCC SIRV Set3 reference with -q flag to ignore the conversion rate calculation for SLAM-seq. To normalize the CPM value for each transcript in each sample based on spike-in RNA, the normalize.loess function of the affy v1.50.0 R package76 was used. Locally estimated scatterplot smoothing (LOESS) normalization was applied to all CPM values of transcripts with the model specified by the distribution of the SIRV-Set-3 spike-ins. Unexpressed genes were filtered out with the cutoff of CPM < 3 in all 8-h and 24-h DMSO and 500 nM dTAG-13 treated samples. ERCC-normalized CPMs of all genes in each sample can be found in **Supplementary Table S2**.

### Assay for Transposase-Accessible Chromatin with high-throughput sequencing (ATAC-seq)

50,000 treated BE(2)-C cells for each sample were washed with 50 μL cold PBS once. Pellets were resuspended in 50 μL cold Omni-ATAC lysis buffer (10 mM Tris-HCl, pH 7.5, 10 mM NaCl, 3 mM MgCl_2_, 0.1% NP-40, 0.1% Tween20, 0.01% digitonin) and incubated on ice for 3 mins, then mixed with 1 mL Omni-ATAC wash buffer (10 mM Tris-HCl, pH 7.5, 10 mM NaCl, 3 mM MgCl2, 0.1% Tween20). Nuclei were pelleted by centrifugation and resuspended in 50 μL Omni-ATAC transposition reaction mixture (25 μL Nextera TD buffer, 16.5 μL PBS, 0.1% Tween20, 0.01% digitonin, 2.5 μL Tn5 transposase) and incubated at 37 °C for 30 mins. DNA was purified using Quick-DNA MicroPrep column purification kit (Zymo Research, #D3020). The libraries were generated by PCR amplification with the cycle numbers determined by qPCR side-reactions as previously described^69^. DNA libraries were purified with Quick-DNA MicroPrep column purification kit (Zymo Research, #D3020). The quality of the libraries was assessed with TapeStation and the DNA contents were quantified by Qubit. The libraries were sequenced on Illumina NextSeq 500 (paired-end, 36-bp reads). Reads were processed with the ENCODE ATAC-seq pipeline (v1.8.0) and were aligned to the human genome build hg19. Adapters were trimmed with Cutadapt (v3.0) and reads were aligned with Bowtie2 (v2.4.2). Blacklist regions excluded using data file wgEncodeDacMapabilityConsensusExcludable.bed.gz. Unmapped/low quality reads were filtered with Samtools (v1.11) and duplicated reads were removed with Picard Tools (v2.23.9). Peaks were called by MACS2 (v2.2.7.1) with call-peak *P* value threshold of 0.01, maximum peaks of 5×10^5^, size of smoothing window of 150, and IDR (irreproducible discovery rate) threshold of 0.05. GC bias computation was performed and TSS (transcription start site) enrichment was calculated on filtered reads. For the differential analysis, the bed files from the same time point were merged with Bedtools (v2.27.1) to generate a peak list including peaks from all treatments. In the merged bed files the adjacent peaks with distances less than 1 kb were merged. The read density calculator Bamliquidator (v1.0) (http://github.com/BradnerLab/pipeline/wiki/bamliquidator) was used to quantify the read density of each treatment at each peak in the corresponding merged bed file with RPM normalization. Peaks with low density (< 0.1 RPM/bp) from all treatments were filtered out. The Log2-transformed fold change and the *P* values given by Student’s *t* test were calculated for each peak.

### Quantitative proteomics

BE(2)C-HDAC2-dTAG cells were treated with 500 nM dTAG-13 for 2 h or 24 h, collected by trypsinization, and washed twice with PBS before flash freezing in liquid nitrogen. Cell pellets were lysed by probe sonication in resuspension buffer (10 mM NaHPO_4_ pH 7.5, 125 mM NaCl, 25 mM KCl, 1.5 mM MgCl_2_, 10% glycerol, 1X HALT Halt Protease Inhibitor Cocktail). Protein concentration was adjusted to 2 mg/mL and 200 μg protein per sample was combined with 48 mg urea to give 8 M final concentration. Samples were reduced by addition of DTT (10 mM final concentration) at 65 °C for 20 mins, before alkylation with iodoacetamide (20 mM) for 30 mins on a 37°C shaking incubator protected from light. Ice-cold H_2_O (500 μL), MeOH (600 μL), and CHCl_3_ (100 μL), were added and the mixture was vortexed and centrifuged (10,000 *g*, 10 mins, 4°C) to afford a protein precipitate at the interface of CHCl_3_ and aqueous layers. Solvent was removed and protein washed with additional MeOH (600 μL) before being allowed to dry briefly at room temperature. The resulting protein pellets were resuspended in EPPS buffer (160 μL, 200 mM, pH 8) by probe sonication. LysC solution (0.5 ug/sample in diH_2_O) was added and the samples were incubated at 37 °C with shaking for 2 h. Trypsin (1 ug in trypsin buffer) and CaCl_2_ (1 μL, 100 mM in H_2_O) were then added and the samples were incubated at 37 °C with shaking overnight. Peptide concentration was determined using the microBCA assay (Thermo Scientific) according to manufacturer’s instructions and samples normalized. To a volume corresponding to 25 μg per sample, CH3CN was added to a final concentration of 30% before incubation with the corresponding TMT tags (60 μg/sample) at room temperature for 30 mins. Additional TMT tag (60 μg/sample) was added and the samples were incubated for another 30 mins. Labeling was quenched by the addition of hydroxylamine (6 μL, 5% in H_2_O). Following a 15 mins incubation at room temperature, formic acid was added to a concentration of 5% and the samples were stored at −80°C until further analysis. Sep Pak desalting, HPLC high-pH fractionation, Orbitrap Fusion (Thermo) Nano-LCMS data collection, and Integrated Proteomics Pipeline (IP2) data analysis were performed as described previously^51^. Data was filtered for 10,000 reporter ion control sum intensity, maximum control channel coefficient of variation < 0.5, and 2 unique peptides identified per protein. Processed data of each sample can be found in **Supplementary Table S3** (2 h) and **Supplementary Table S4** (24 h).

### Gene set enrichment analysis (GSEA)

The averages of the replicates from the control group or the treatment group were calculated respectively for each gene and used as the input for GSEA. The GSEA software developed by UC San Diego and Broad Institute^70,71^ was used with the following parameters: gene sets database c2.all, number of permutations 1000, collapse no_collapse, permutation type gene_set.

### qRT-PCR

RNA was extracted with the RNeasy miniprep kit (Qiagen). 2.5 μg of RNA from each sample was used to synthesize cDNA with the SuperScript VILO cDNA Synthesis Kit (Life Technologies, #11755050). Quantification of each transcript of interest was performed in triplicates with SYBR Select Master Mix (Life Technologies, #4472908) on BioRad CFX 384 with primer pairs listed below. Cycle threshold (Ct) values were determined by the default setting on BioRab CFX Manager (v3.0) and the relative expression level of each gene was calculated with normalization to *B2M* transcript levels and DMSO control using the ddCt method.

### Primer pairs for qRT-PCR

*B2M*: forward 5’-TCTCTGCTGGATGACGTGAG-3’, reverse 5’-TAGCTGTGCTCGCGCTACT-3’. *GAPDH*: forward 5’-CATCATCCCTGCCTCTACTG-3’, reverse 5’-GCCTGCTTCACCACCTTC-3’. *ACTB*: forward 5’-TGGCACCCAGCACAATGAA-3’, reverse 5’-CTAAGTCATAGTCCGCCTAGAAGCA-3’. *CHD4*: forward 5’-GCTGCAACCATCCATACCTC-3’, reverse 5’-ACCATCGATGCGTTCGTATT-3’. *HDAC1:* forward 5’-GGAAATCTATCGCCCTCACA-3’, reverse 5’-AACAGGCCATCGAATACTGG-3’. *GATAD2A*: forward 5’-ACGAGTTCATCTACCTGGTCGG-3’, reverse 5’-ACGTGAAGTCCGTCTTGCACTG-3’. *GATAD2B*: forward 5’-CAAAAGCTGTGCCTCACTTC-3’, reverse 5’-TTCCAGTGAGGGGTGAAATC-3’. *MBD2*: forward 5’-AAGTGCTGGCAAGAGCGATGTCTA-3’, reverse 5’-TTTCCCAGGTACCTTGCCAACTGA-3’. *MBD3*: forward 5’-GGCCACAGGGATGTCTTTTACT-3’, reverse 5’-CGGCTTGCTGCGGAACT-3’. *MTA1:* forward 5’-TGCTCAACGGGAAGTCCTACC-3’, reverse 5’-GGGCATGTAGAACACGTCACC-3’.*MTA2*: forward 5’-TATCACTCTGTTTCACGCCA-3’, reverse 5’-ACCATTCCTCCATCTCATCC-3’. *MTA3*: forward 5’-AAGCCTGGTGCTGTGAAT-3’, reverse 5’-AGGGTCCTCTGTAGTTGG-3’. *IRS2:* forward 5’-GAGTGCACCCGTACCTATGGAA-3’, reverse 5’-GAAATCCGGCTTTACCTTGAACT-3’.

## Acknowledgements

We gratefully acknowledge Dr. Martin G. Jaeger for his critically reading of the manuscript. This work was supported by the National Institutes of Health (NIH) through an NIH Director’s Early Independence Award (DP5-OD26380) to M.A.E. and by the Ono Pharma Foundation to M.A.E..

## Extended data figures

**Extended Data Fig. 1.**
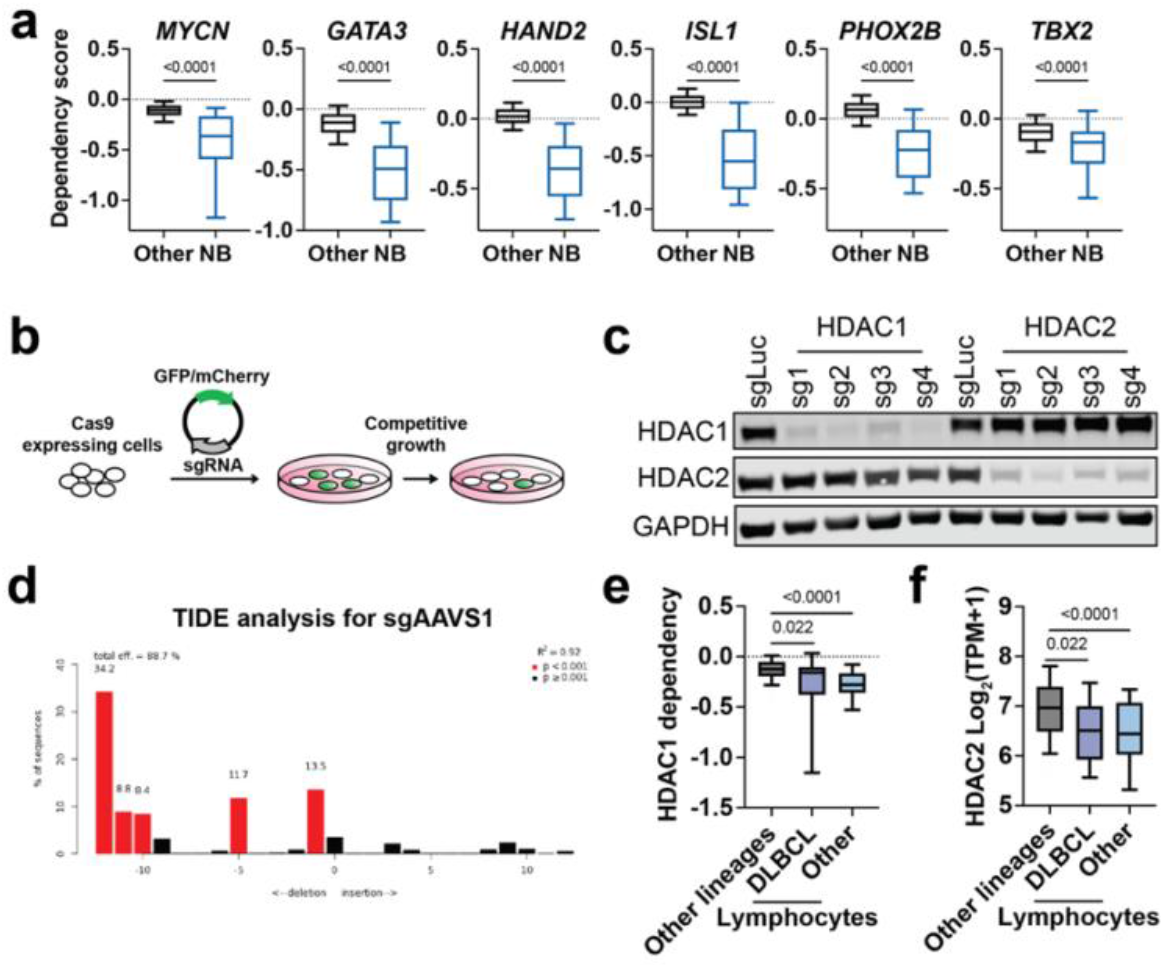
*HDAC1* and *HDAC2* are selective dependencies in neuroblastoma and lymphoid malignancies. **a**, Boxplots of dependency scores of core regulatory circuitry (CRC) transcription factor genes in neuroblastoma. **b**, Schematic illustration of CRISPR-Cas9-based competitive growth assay. **c**, Validation of on-target effects of *HDAC1* and *HDAC2* guides by immunoblot. **d**, TIDE (tracking of indels by decomposition) analysis shows a cutting efficiency of 88.7% for sgAAVS1. **e,** Boxplots of *HDAC1* dependnecy scores in DLBCL lines (purple, *n* = 8), non-DLBCL lymphocyte lines (blue, n = 19), and other lineages (grey, *n* = 962). **f**, Boxplot of *HDAC2* transcript levels in DLBCL lines (purple, *n* = 20), non-DLBCL lymphocyte lines (blue, *n* = 63), and other lineages (grey, *n* = 1,292). Data from DepMap, CRISPR_genetic_effect 22Q1 and CCLE_expression 22Q1. *P* values were obtained via two-tailed Student’s *t*-test. Boxplots represent 25-75 percentiles with whiskers extending to 10-90 percentiles.

**Extended Data Fig. 2.**
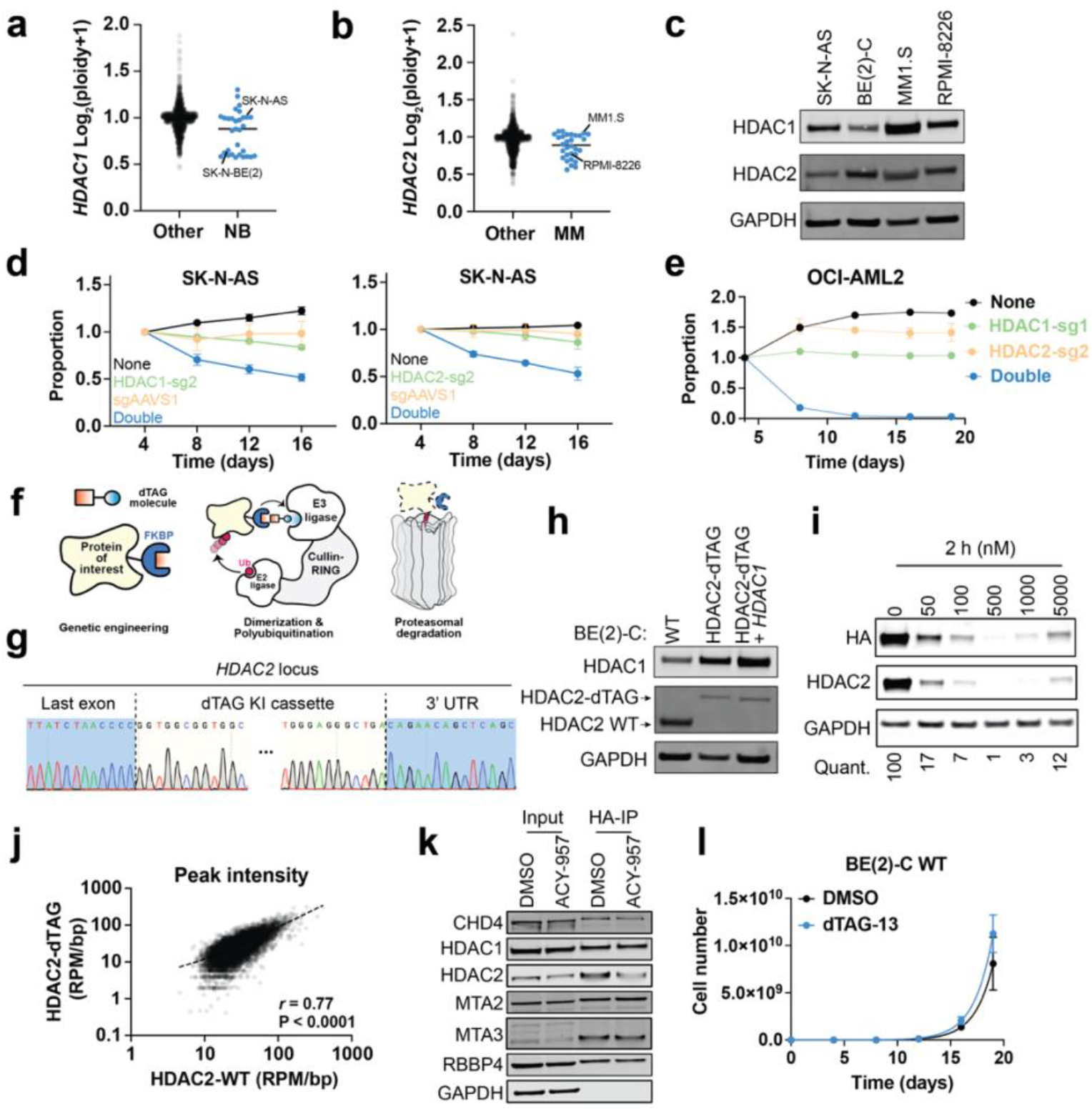
dTAG tagging does not affect HDAC2 normal function and allows efficient degradation upon dTAG-13 treatment. **a**, *HDAC1* copy number in neuroblastoma (*n* = 31) and other cell lines (*n* = 1,340). **b**, *HDAC2* copy number in multiple myeloma (*n* = 30) and other cell lines (*n* = 1,341). **c**, *HDAC1* and *HDAC2* expression levels determined by immunoblot. **d**, Control groups related to the two-color competitive growth assay in (**Fig. 2c**). **e**, Two-color competitive growth assay with HDAC1-sg1 and HDAC2-sg2 in OCI-AML2 cells. **f**, Schematic illustration of the dTAG system. dTAG PROTACs mediate dimerization of the FKBP12^F36A^-tagged protein of interest and an E3 ubiquitin ligase, which results in ubiquitination and proteasomal degradation of the target protein. **g**, Representative Sanger sequencing chromatograms of *HDAC2* locus of a clone with successful dTAG knock-in. **h**, Immunoblot validation of HDAC2-dTAG cell lines with and without *HDAC1* overexpression. **i**, Dose response of dTAG-13 treatment in BE(2)-C-HDAC2-dTAG cells (2 h). **j**, Correlation of HDAC2 CUT&RUN in BE(2)-C cells versus HA CUT&RUN in BE(2)-C-HDAC2-dTAG cells. *P* value was determined by Pearson correlation coefficient (*r*). *n* = 10,832. **k**, Co-immunoprecipitation of HDAC2-dTAG (IP: HA) and NuRD subunits. **l**, Proliferation of BE(2)-C wild-type cells are not affected by dTAG-13 (500 nM) (blue) compared to the DMSO group (black).

**Extended Data Fig. 3.**
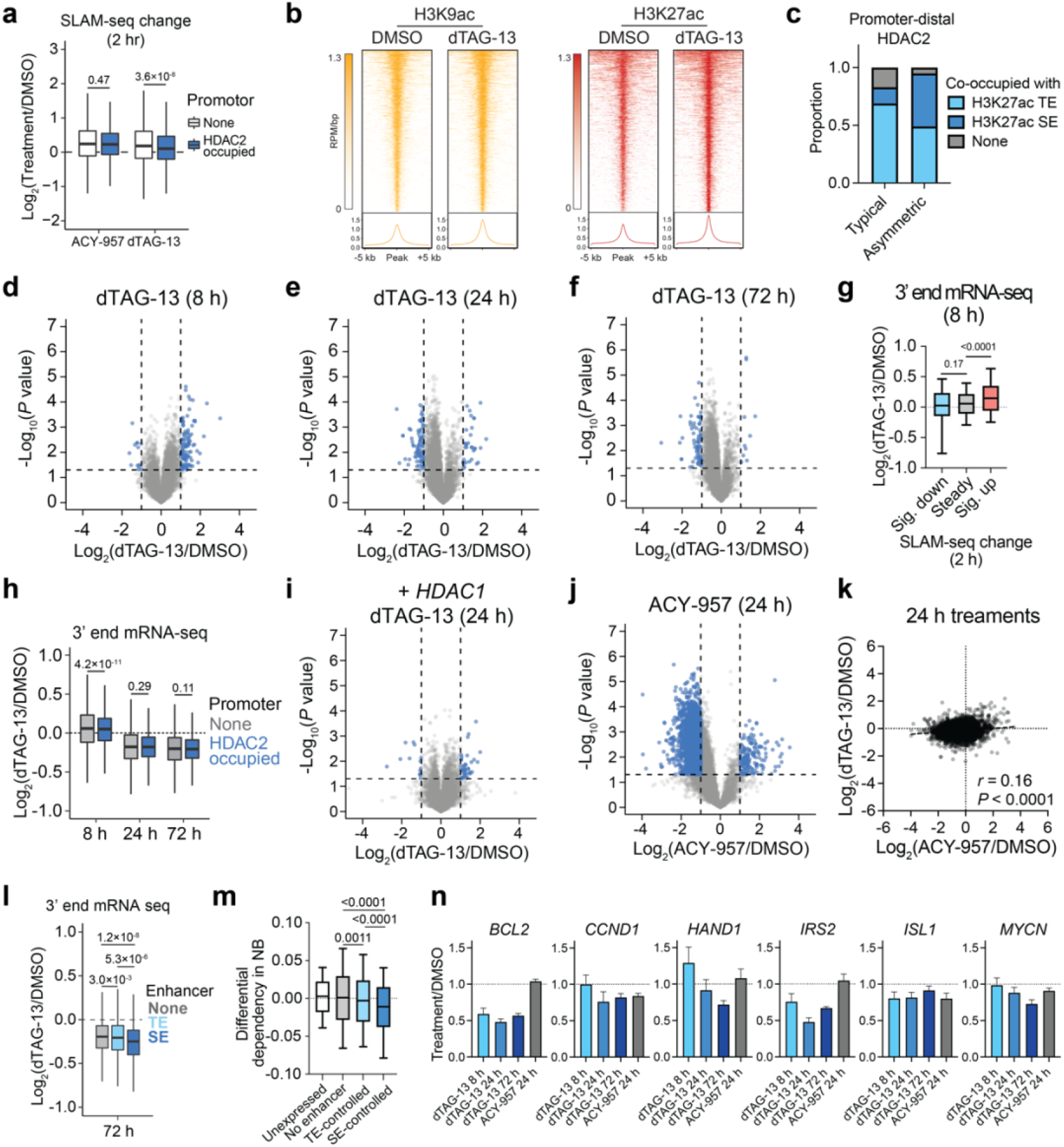
HDAC2 degradation disrupts transcriptional regulation. **a**, Boxplot of SLAM-seq changes at genes with (*n* = 5,082) or without (*n* = 8,117) HDAC2 bound at the promoter (TSS ± 1kb) with ACY-957 (5 μM) or dTAG-13 (500 nM) for 2 h. **b**, Heatmaps (top) and metaplots (bottom) of CUT&RUN signals with DMSO control and 2-h dTAG-13 (500 nM) treatment. **c**, Genomic feature distribution of typical and asymmetric HDAC2 sites shows a biased occupancy of super enhancers by asymmetric sites. **d-f**, Volcano plots of changes in total mRNA abundance (3’-end mRNA-seq) following dTAG-13 treatment (500 nM) for 8 h (**d**), 24 h (**e**), and 72 h (**f**). *n* = 3. *P* values were determined by two-tailed Student’s *t*-test. **g**, Boxplots of total mRNA transcript changes following 8-h dTAG-13 (500 nM) treatment. Genes are grouped by 2-h SLAM-seq data (shown in **Fig. 3a**): significantly downregulated (Log2(dTAG-13/DMSO) < −1 and *P* value < 0.05, *n* = 59), significantly upregulated (Log_2_(dTAG-13/DMSO) > 1 and *P* value < 0.05, *n* = 224), and unchanged (all other genes, *n* = 12,916). Boxes represent 25-75 percentiles with whiskers extending to 10-90 percentiles. **h**, Boxplots of total transcript change of genes with (*n* = 5,082) or without (*n* = 8,117) HDAC2 bound at the promoter. **i,j**Volcano plot of total transcript changes following 24-h dTAG-13 (500 nM) treatment in BE(2)-C-HDAC2-dTAG cells overexpressing *HDAC1* (**i**) or ACY-957 treatment (5 μM) in BE(2)-C-HDAC2-dTAG cells (**j**). *P* values were determined by two-tailed Student’s *t*-test. **k**, Correlation of total transcript changes (3’-end mRNA-seq) following 24-h treatment with dTAG-13 (500 nM) or ACY-957 (5 μM) in BE(2)-C-HDAC2-dTAG cells. *P* value was determined by Pearson correlation coefficient (*r*). *n* = 13,199. **l**, Boxplot of total mRNA transcript changes following 72-h dTAG-13 treatment (500 nM) for genes not associated with an enhancer (*n* = 7,588), associated with typical enhancers (*n* = 5,026), or associated with super enhancers (*n* = 585) in BE(2)C-HDAC2-dTAG cells. **m**, Boxplot of neuroblastoma-specific differential dependencies for genes that in BE(2)-C cells are unexpressed (CPM < 3 shown by 3’-end mRNA-seq data, *n* = 7,973), expressed but not associated with an enhancer (*n* = 7,588), expressed and associated with a typical enhancer (*n* = 5,026), or expressed and associated with a super enhancers (*n* = 585). Boxes represent 25-75 percentiles with whiskers extending to 10-90 percentiles. **n**, Bar plots of drug-induced mRNA transcript changes for select neuroblastoma-specific dependencies. Normalized to DMSO. Mean ± s.d., *n* = 3. Unless specified, *P* values were determined by two-tailed Welch’s *t*-test. Unless specified, boxplots represent 25-75 percentiles with whiskers extending 1.5 IQR.

**Extended Data Fig. 4.**
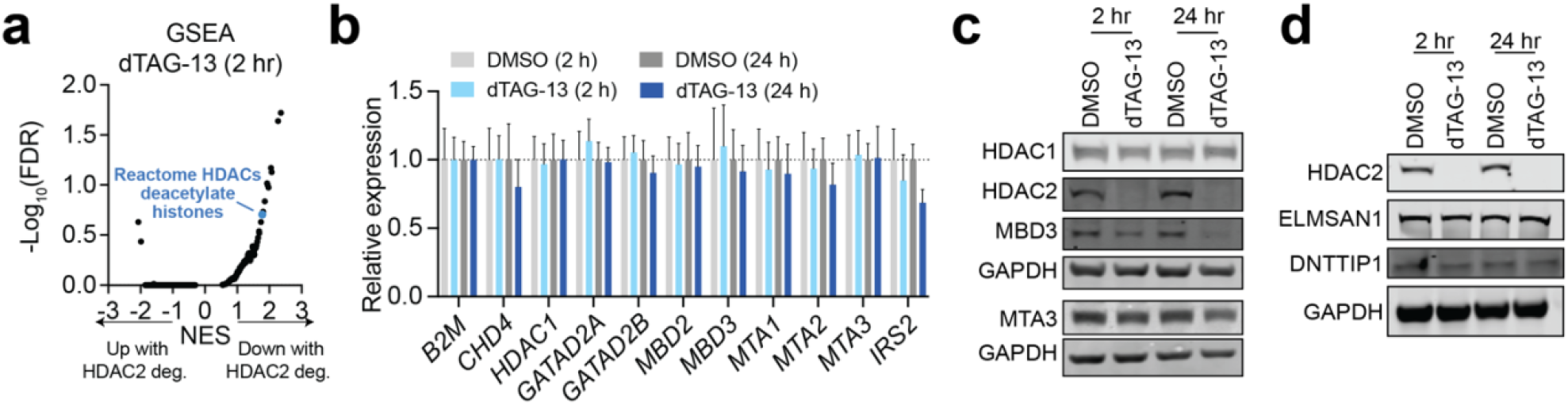
HDAC2 loss destabilizes the NuRD complex. **a**, GSEA of proteomic changes following 2-h dTAG-13 (500 nM) treatment. **b**, DMSO-normalized changes in gene expression (qRT-PCR) following dTAG-13 treatments (500 nM). Gene expression levels are normalized to *B2M* transcript level, *n* = 3. *IRS2* was used as a positive control as it was significantly downregulated by 24-h dTAG-13 treatment shown by 3’-end mRNA-seq (**Extended Data Fig.3n**). **c**,**d** Immunoblots of NuRD subunits (**d**) and MiDAC subunits (**e**) with 2-h and 24-h dTAG-13 treatments in BE(2)-C-HDAC2-dTAG cells.

**Extended Data Fig. 5.**
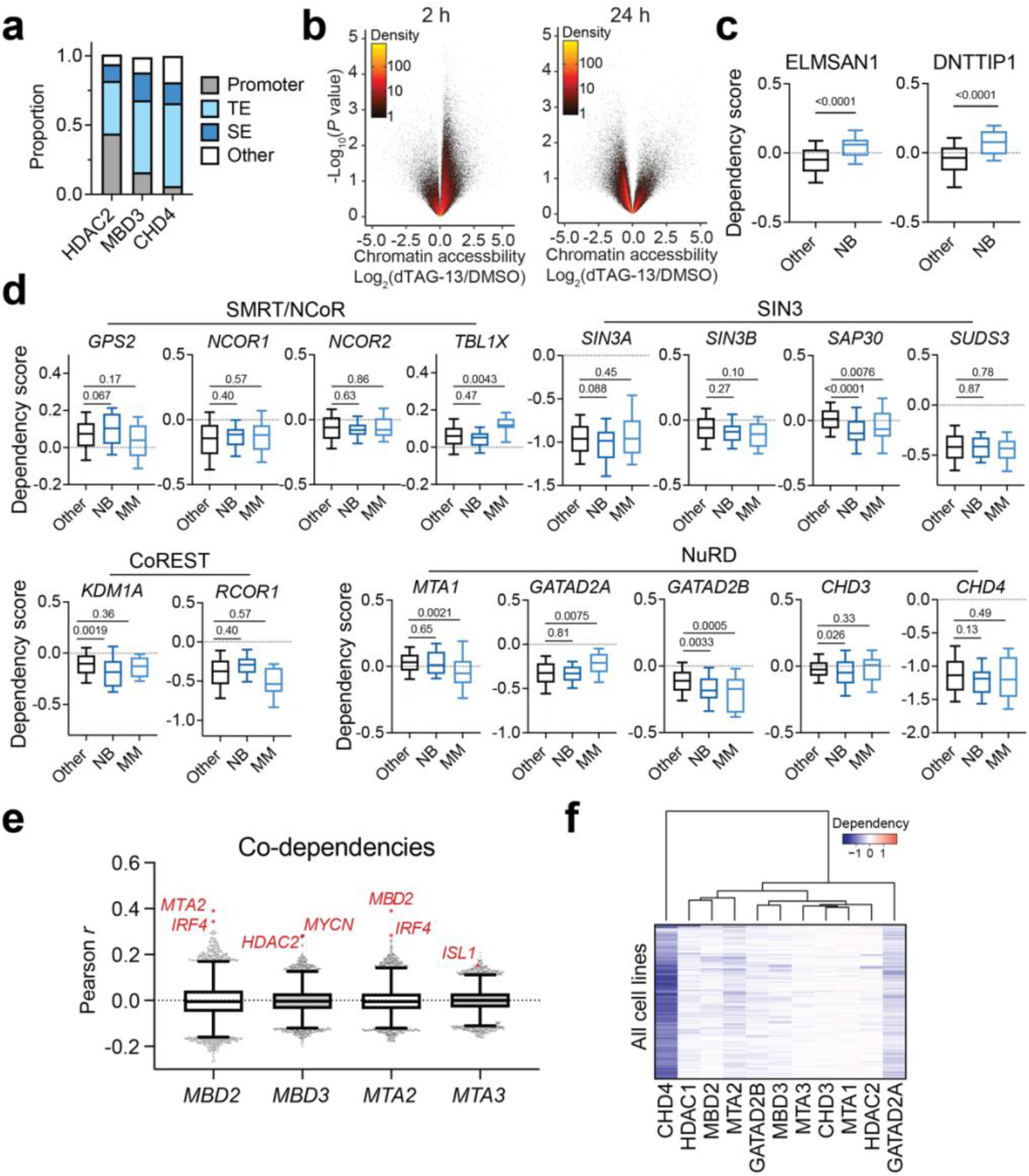
Destabilized NuRD complex leads to dysregulated chromatin accessibility. **a**, Genomic feature distribution of sites bound by HDAC2, MBD3, and CHD4. **b,** Volcano plots of chromatin accessibility changes measured by ATAC-seq following dTAG-13 (500 nM) treatment for 2 h (left) or 24 h (right). *n* = 3. **c**, Dependency scores of MiDAC subunits in neuroblastoma (*n* = 34) and other cell lines (*n* = 1,020). **d**, Dependency scores of HDAC1/2-containing complexes in neuroblastoma (*n* = 34), multiple myeloma (*n* = 21), and other cell lines (*n* = 999). **e**, Boxplots of Pearson correlation coefficients between *MBD2, MBD3, MTA2,* and *MTA3* dependency scores compared to all other genes (*n* = 17,386). **f**, Unsupervised clustering of dependency scores of NuRD subunits in all cell lines. *P* values were obtained via two-tailed Student’s *t*-test. Boxplots represent 25-75 percentiles with whiskers extending to 10-90 percentiles.

## References

1. Shortt J, Ott CJ, Johnstone RW, Bradner JE. A chemical probe toolbox for dissecting the cancer epigenome. Nat Rev Cancer. 2017;17(3):160–183. doi:10.1038/nrc.2016.148

2. Chang L, Ruiz P, Ito T, Sellers WR. Targeting pan-essential genes in cancer: Challenges and opportunities. Cancer Cell. 2021;39(4):466–479. doi:10.1016/j.ccell.2020.12.008

3. Seto E, Yoshida M. Erasers of Histone Acetylation: The Histone Deacetylase Enzymes. Cold Spring Harb Perspect Biol. 2014;6(4):a018713–a018713. doi:10.1101/cshperspect.a018713

4. Taunton J, Hassig CA, Schreiber SL. A Mammalian Histone Deacetylase Related to the Yeast Transciptional Regulator Rpd3p Published by: American Association for the Advancement of Science Stable URL: https://www.jstor.org/stable/2890320 American Association for the Advancement of Science is. Science (80-). 1996;272(5260):408–411.

5. Millard CJ, Watson PJ, Fairall L, Schwabe JWR. Targeting Class I Histone Deacetylases in a “Complex” Environment. Trends Pharmacol Sci. 2017;38(4):363–377. doi:10.1016/j.tips.2016.12.006

6. Li Y, Seto E. HDACs and HDAC inhibitors in cancer development and therapy. Cold Spring Harb Perspect Med. 2016;6(10):1–34. doi:10.1101/cshperspect.a026831

7. Ito T, Young MJ, Li R, et al. Paralog knockout profiling identifies DUSP4 and DUSP6 as a digenic dependence in MAPK pathway-driven cancers. Nat Genet. 2021;53(12):1664–1672. doi:10.1038/s41588-021-00967-z

8. DeWeirdt PC, Sanson KR, Sangree AK, et al. Optimization of AsCas12a for combinatorial genetic screens in human cells. Nat Biotechnol. 2021;39(1):94–104. doi:10.1038/s41587-020-0600-6

9. Huang A, Garraway LA, Ashworth A, Weber B. Synthetic lethality as an engine for cancer drug target discovery. Nat Rev Drug Discov. 2020;19(1):23–38. doi:10.1038/s41573-019-0046-z

10. Bryant HE, Schultz N, Thomas HD, et al. Specific killing of BRCA2-deficient tumours with inhibitors of poly(ADP-ribose) polymerase.[erratum appears in Nature. 2007 May 17;447(7142):346]. Nature. 2005;434(7035):913–917.

11. Hoffman GR, Rahal R, Buxton F, et al. Functional epigenetics approach identifies BRM/SMARCA2 as a critical synthetic lethal target in BRG1-deficient cancers. Proc Natl Acad Sci U S A. 2014;111(8):3128–3133. doi:10.1073/pnas.1316793111

12. Wilson BG, Helming KC, Wang X, et al. Residual Complexes Containing SMARCA2 (BRM) Underlie the Oncogenic Drive of SMARCA4 (BRG1) Mutation. Mol Cell Biol. 2014;34(6):1136–1144. doi:10.1128/mcb.01372-13

13. Oike T, Ogiwara H, Tominaga Y, et al. A synthetic lethality-based strategy to treat cancers harboring a genetic deficiency in the chromatin remodeling factor BrG1. Cancer Res. 2013;73(17):5508–5518. doi:10.1158/0008-5472.CAN-12-4593

14. Ogiwara H, Sasaki M, Mitachi T, et al. Targeting p300 addiction in CBP-deficient cancers causes synthetic lethality by apoptotic cell death due to abrogation of MYC expression. Cancer Discov. 2016;6(4):430–445. doi:10.1158/2159-8290.CD-15-0754

15. Lelij P van der, Lieb S, Jude J, et al. Synthetic lethality between the cohesin subunits STAG1 and STAG2 in diverse cancer contexts. Elife. 2017;6:e26980. doi:10.7554/elife.26980

16. Tsherniak A, Vazquez F, Montgomery PG, et al. Defining a Cancer Dependency Map. Cell. 2017;170(3):564–576.e16. doi:10.1016/j.cell.2017.06.010

17. Meyers RM, Bryan JG, McFarland JM, et al. Computational correction of copy number effect improves specificity of CRISPR-Cas9 essentiality screens in cancer cells. Nat Genet. 2017;49(12):1779–1784. doi:10.1038/ng.3984

18. Parrish PCR, Thomas JD, Gabel AM, Kamlapurkar S, Bradley RK, Berger AH. Discovery of synthetic lethal and tumor suppressor paralog pairs in the human genome. Cell Rep. 2021;36(9):109597. doi:10.1016/j.celrep.2021.109597

19. Malone CF, Dharia N V., Kugener G, et al. Selective modulation of a pan-essential protein as a therapeutic strategy in cancer. Cancer Discov. 2021;11(9):2282–2299. doi:10.1158/2159-8290.CD-20-1213

20. Caron H, van Sluis P, van Hoeve M, et al. Allelic loss of chromosome 1p36 in neuroblastoma is of preferential maternal origin and correlates with N–myc amplification. Nat Genet. 1993;4(2):187–190. doi:10.1038/ng0693-187

21. Maris JM, White PS, Beltinger CP, et al. Significance of Chromosome 1p Loss of Heterozygosity in Neuroblastoma. Cancer Res. 1995;55(20):4664–4669.

22. Janoueix-Lerosey I, Novikov E, Monteiro M, et al. Gene expression profiling of 1 p35-36 genes in neuroblastoma. Oncogene. 2004;23(35):5912–5922. doi:10.1038/sj.onc.1207784

23. Komotar RJ, Otten ML, Starke RM, Anderson RCE. Chromosome 1p and 11q deletions and outcome in neuroblastoma—A critical review. Clin Med Oncol. 2008;2:419–420. doi:10.4137/cmo.s391

24. Merup M, Moreno TC, Heyman M, et al. 6q deletions in acute lymphoblastic leukemia and non-Hodgkin’s lymphomas. Blood. 1998;91(9):3397–3400. doi:10.1182/blood.v91.9.3397

25. Thelander EF, Ichimura K, Corcoran M, et al. Characterization of 6q deletions in mature B cell lymphomas and childhood acute lymphoblastic leukemia. Leuk Lymphoma. 2008;49(3):477–487. doi:10.1080/10428190701817282

26. Taborelli M, Tibiletti MG, Martin V, Pozzi B, Bertoni F, Capella C. Chromosome band 6q deletion pattern in malignant lymphomas. Cancer Genet Cytogenet. 2006;165(2):106–113. doi:10.1016/j.cancergencyto.2005.06.025

27. Aktas Samur A, Minvielle S, Shammas M, et al. Deciphering the chronology of copy number alterations in Multiple Myeloma. Blood Cancer J. 2019;9(4). doi:10.1038/s41408-019-0199-3

28. Durbin AD, Zimmerman MW, Dharia N V., et al. Selective gene dependencies in MYCN-amplified neuroblastoma include the core transcriptional regulatory circuitry. Nat Genet. 2018;50(9):1240–1246. doi:10.1038/s41588-018-0191-z

29. Zeid R, Lawlor MA, Poon E, et al. Enhancer invasion shapes MYCN-dependent transcriptional amplification in neuroblastoma. Nat Genet. 2018;50(4):515–523. doi:10.1038/s41588-018-0044-9

30. Dharia N V., Kugener G, Guenther LM, et al. A first-generation pediatric cancer dependency map. Nat Genet. 2021;53(4):529–538. doi:10.1038/s41588-021-00819-w

31. Chen L, Alexe G, Dharia N V., et al. CRISPR-Cas9 screen reveals a MYCN-amplified neuroblastoma dependency on EZH2. J Clin Invest. 2018;128(1):446–462. doi:10.1172/JCI90793

32. Brinkman EK, Chen T, Amendola M, Van Steensel B. Easy quantitative assessment of genome editing by sequence trace decomposition. Nucleic Acids Res. 2014;42(22):1–8. doi:10.1093/nar/gku936

33. Ghandi M, Huang FW, Jané-Valbuena J, et al. Next-generation characterization of the Cancer Cell Line Encyclopedia. Nature. 2019;569(7757):503–508. doi:10.1038/s41586-019-1186-3

34. Cerami E, Gao J, Dogrusoz U, et al. The cBio Cancer Genomics Portal: An open platform for exploring multidimensional cancer genomics data. Cancer Discov. 2012;2(5):401–404. doi:10.1158/2159-8290.CD-12-0095

35. Frumm SM, Fan ZP, Ross KN, et al. Selective HDAC1/HDAC2 inhibitors induce neuroblastoma differentiation. Chem Biol. 2013;20(5):713–725. doi:10.1016/j.chembiol.2013.03.020

36. Jaeger MG, Winter GE. Fast-acting chemical tools to delineate causality in transcriptional control. Mol Cell. 2021;81(8):1617–1630. doi:10.1016/j.molcel.2021.02.015

37. Zhang Y, Erb MA. Enabling cancer target validation with genetically encoded systems for ligand-induced protein degradation. Curr Res Chem Biol. 2021;1:100011. doi:10.1016/j.crchbi.2021.100011

38. Erb MA, Scott TG, Li BE, et al. Transcription control by the ENL YEATS domain in acute leukaemia. Nature. 2017;543(7644):270–274. doi:10.1038/nature21688

39. Nabet B, Roberts JM, Buckley DL, et al. The dTAG system for immediate and target-specific protein degradation. Nat Chem Biol. 2018;14(5):431–441. doi:10.1038/s41589-018-0021-8

40. Skene PJ, Henikoff JG, Henikoff S. Targeted in situ genome-wide profiling with high efficiency for low cell numbers. Nat Protoc. 2018;13(5):1006–1019. doi:10.1038/nprot.2018.015

41. Herzog VA, Reichholf B, Neumann T, et al. Thiol-linked alkylation of RNA to assess expression dynamics. Nat Methods. 2017;14(12):1198–1204. doi:10.1038/nmeth.4435

42. Shearstone JR, Golonzhka O, Chonkar A, et al. Chemical inhibition of histone deacetylases 1 and 2 induces fetal hemoglobin through activation of GATA2. PLoS One. 2016;11(4):1–27. doi:10.1371/journal.pone.0153767

43. Whyte WA, Orlando DA, Hnisz D, et al. Master transcription factors and mediator establish super-enhancers at key cell identity genes. Cell. 2013;153(2):307–319. doi:10.1016/j.cell.2013.03.035

44. Schölz C, Weinert BT, Wagner SA, et al. Acetylation site specificities of lysine deacetylase inhibitors in human cells. Nat Biotechnol. 2015;33(4):415–425. doi:10.1038/nbt.3130

45. Marques JG, Gryder BE, Pavlovic B, et al. NURD subunit CHD4 regulates super-enhancer accessibility in rhabdomyosarcoma and represents a general tumor dependency. Elife. 2020;9:1–30. doi:10.7554/ELIFE.54993

46. Gryder BE, Pomella S, Sayers C, et al. Histone hyperacetylation disrupts core gene regulatory architecture in rhabdomyosarcoma. Nat Genet. 2019;51(12):1714–1722. doi:10.1038/s41588-019-0534-4

47. Xiong Y, Donovan KA, Eleuteri NA, et al. Chemo-proteomics exploration of HDAC degradability by small molecule degraders. Cell Chem Biol. 2021;28(10):1514–1527.e4. doi:10.1016/j.chembiol.2021.07.002

48. Hsu JHR, Rasmusson T, Robinson J, et al. EED-Targeted PROTACs Degrade EED, EZH2, and SUZ12 in the PRC2 Complex. Cell Chem Biol. 2020;27(1):41–46.e17. doi:10.1016/j.chembiol.2019.11.004

49. Farnaby W, Koegl M, Roy MJ, et al. BAF complex vulnerabilities in cancer demonstrated via structure-based PROTAC design. Nat Chem Biol. 2019;15(7):672–680. doi:10.1038/s41589-019-0294-6

50. Schick S, Grosche S, Kohl KE, et al. Acute BAF perturbation causes immediate changes in chromatin accessibility. Nat Genet. 2021;53(3):269–278. doi:10.1038/s41588-021-00777-3

51. Vinogradova E V., Zhang X, Remillard D, et al. An Activity-Guided Map of Electrophile-Cysteine Interactions in Primary Human T Cells. Cell. 2020;182(4):1009–1026.e29. doi:10.1016/j.cell.2020.07.001

52. Denslow SA, Wade PA. The human Mi-2/NuRD complex and gene regulation. Oncogene. 2007;26(37):5433–5438. doi:10.1038/sj.onc.1210611

53. Low JKK, Silva APG, Sharifi Tabar M, et al. The Nucleosome Remodeling and Deacetylase Complex Has an Asymmetric, Dynamic, and Modular Architecture. Cell Rep 2020; 33(9):108450. doi:10.1016/j.celrep.2020.108450

54. Millard CJ, Varma N, Saleh A, et al. The structure of the core NuRD repression complex provides insights into its interaction with chromatin. Elife. 2016;5:1–21. doi:10.7554/elife.13941

55. Sher F, Hossain M, Seruggia D, et al. Rational targeting of a NuRD subcomplex guided by comprehensive in situ mutagenesis. Nat Genet. 2019;51(7):1149–1159. doi:10.1038/s41588-019-0453-4

56. Buenrostro JD, Wu B, Chang HY, Greenleaf WJ. ATAC-seq: A method for assaying chromatin accessibility genome-wide. Curr Protoc Mol Biol. 2015;2015(January):21.29.1–21.29.9. doi:10.1002/0471142727.mb2129s109

57. Gryder BE, Wu L, Woldemichael GM, et al. Chemical genomics reveals histone deacetylases are required for core regulatory transcription. Nat Commun. 2019;10(1):1–12. doi:10.1038/s41467-019-11046-7

58. Qu K, Zaba LC, Satpathy AT, et al. Chromatin Accessibility Landscape of Cutaneous T Cell Lymphoma and Dynamic Response to HDAC Inhibitors. Cancer Cell. 2017;32(1):27–41.e4. doi:10.1016/j.ccell.2017.05.008

59. Pan J, Meyers RM, Michel BC, et al. Interrogation of Mammalian Protein Complex Structure, Function, and Membership Using Genome-Scale Fitness Screens. Cell Syst. 2018;6(5):555–568.e7. doi:10.1016/j.cels.2018.04.011

60. Michel BC, D’Avino AR, Cassel SH, et al. A non-canonical SWI/SNF complex is a synthetic lethal target in cancers driven by BAF complex perturbation. Nat Cell Biol. 2018;20(12):1410–1420. doi:10.1038/s41556-018-0221-1

61. Lai AY, Wade PA. Cancer biology and NuRD: A multifaceted chromatin remodelling complex. Nat Rev Cancer. 2011;11(8):588–596. doi:10.1038/nrc3091

62. Bornelöv S, Reynolds N, Xenophontos M, et al. The Nucleosome Remodeling and Deacetylation Complex Modulates Chromatin Structure at Sites of Active Transcription to Fine-Tune Gene Expression. Mol Cell. 2018;71(1):56–72.e4. doi:10.1016/j.molcel.2018.06.003

63. Smalley JP, Baker IM, Pytel WA, et al. Optimization of Class I Histone Deacetylase PROTACs Reveals that HDAC1/2 Degradation is Critical to Induce Apoptosis and Cell Arrest in Cancer Cells. J Med Chem. Published online 2022. doi:10.1021/acs.jmedchem.1c02179

64. Scholes NS, Mayor-Ruiz C, Winter GE. Identification and selectivity profiling of small-molecule degraders via multi-omics approaches. Cell Chem Biol. 2021;28(7):1048–1060. doi:10.1016/j.chembiol.2021.03.007

65. Sakuma T, Nakade S, Sakane Y, Suzuki KIT, Yamamoto T. MMEJ-Assisted gene knock-in using TALENs and CRISPR-Cas9 with the PITCh systems. Nat Protoc. 2016;11(1):118–133. doi:10.1038/nprot.2015.140

66. Shi J, Wang E, Milazzo JP, Wang Z, Kinney JB, Vakoc CR. Discovery of cancer drug targets by CRISPR-Cas9 screening of protein domains. Nat Biotechnol. 2015;33(6):661–667. doi:10.1038/nbt.3235

67. Doench JG, Fusi N, Sullender M, et al. Optimized sgRNA design to maximize activity and minimize off-target effects of CRISPR-Cas9. Nat Biotechnol. 2016;34(2):184–191. doi:10.1038/nbt.3437

68. Neumann T, Herzog VA, Muhar M, et al. Quantification of experimentally induced nucleotide conversions in high-throughput sequencing datasets. BMC Bioinformatics. 2019;20(1):1–16. doi:10.1186/s12859-019-2849-7

69. Corces MR, Trevino AE, Hamilton EG, et al. An improved ATAC-seq protocol reduces background and enables interrogation of frozen tissues. Nat Methods. 2017;14(10):959–962. doi:10.1038/nmeth.4396

70. Subramanian A, Tamayo P, Mootha VK, et al. Gene set enrichment analysis: A knowledge-based approach for interpreting genome-wide expression profiles. Proc Natl Acad Sci U S A. 2005;102(43):15545–15550. doi:10.1073/pnas.0506580102

71. Mootha VK, Lindgren CM, Eriksson K-F, et al. PGC-1α-responsive genes involved in oxidative phosphorylation are coordinately downregulated in human diabetes. Nat Genet. 2003;34(3):267–273.

